# Quality-controlled R-loop meta-analysis reveals the characteristics of R-Loop consensus regions

**DOI:** 10.1101/2021.11.01.466823

**Authors:** H. E. Miller, D. Montemayor, J. Abdul, A. Vines, S. Levy, S. Hartono, K. Sharma, B. Frost, F. Chedin, A. J. R. Bishop

## Abstract

R-loops are three-stranded nucleic acid structures formed from the hybridization of RNA and DNA during transcription. While the pathological consequences of R-loops have been well-studied to date, the locations, classes, and dynamics of physiological R-loops remain poorly understood. R-loop mapping studies provide insight into R-loop dynamics, but their findings are challenging to generalize. This is due to the narrow biological scope of individual studies, the limitations of each mapping modality, and, in some cases, poor data quality. In this study, we reprocessed 693 R-loop mapping datasets from a wide array of biological conditions and mapping modalities. From this data resource, we developed an accurate method for R-loop data quality control, and we reveal the extent of poor-quality data within previously published studies. We then identified a set of high-confidence R-loop mapping samples and used them to define consensus R-loop sites called “R-loop regions” (RL regions). In the process, we revealed the stark divergence between S9.6 and dRNH-based R-loop mapping methods and identified biologically meaningful subtypes of both constitutive and variable R-loops. Taken together, this work provides a much-needed method to assess R-loop data quality and reveals intriguing aspects of R-loop biology.

## INTRODUCTION

R-loops are three-stranded nucleic acid structures formed from the hybridization of RNA and DNA, often in regions with high G (guanine) or C (cytosine) skew, such as CpG islands (1–3). The study of R-loops has mostly focused on their pathological consequences in promoting genomic instability (4, 5) and in hypertranscriptional cancers, such as Ewing sarcoma (6, 7). However, recent studies have revealed that R-loops contribute to physiological processes, such as DNA repair (8), DNA methylation (9), and even reprogramming to pluripotency (10). While these studies have led to an increase in interest regarding the cause and consequences of physiological R-loops, fundamental questions regarding their subtypes, dynamics, and interactions remain unanswered.

R-loop mapping via high-throughput sequencing can elucidate R-loop locations and dynamics genome-wide. Presented in 2012, DNA:RNA immunoprecipitation (DRIP) sequencing was the first such technique, utilizing the S9.6 antibody to detect R-loops (2). In 2017, *Chen et al*. introduced “R-ChIP”, an R-loop mapping modality which relies on catalytically inactive RNase H1 (dRNH) as an alternative to S9.6-based methods (11). Though preliminary evidence suggested differences between dRNH and S9.6-based R-loop mapping exist (11, 12), the extent and biological relevance of these disparities remains unclear. Recently, *Castillo-Guzman and Chedin* reviewed the relevant R-loop mapping literature and hypothesized that dRNH and S9.6 may map biologically distinct subtypes of R-loops (“Class I and II”) (13), but no studies to date have evaluated these subtypes bioinformatically. It is also unknown how different R-loop mapping modalities relate to each other more generally or whether they each reliably map the same sets of R-loops. This gap in knowledge has become a critical concern as, at the time of writing, there are now 23 distinct R-loop mapping modalities, 14 of which were developed since 2019 alone (**Table S1**).

Previous work in the transcriptomics and epigenomics fields have demonstrated the potential for public data mining to yield novel biological insights (14, 15). However, data mining studies are scant within the R-loop field. Our previous analysis of 108 public DRIP-Seq datasets is the largest such study to date (16). It demonstrated the potential for greater insight from larger and more diverse datasets. However, it also revealed a number of inconsistencies in quality between publicly available R-loop mapping samples (16). These findings were echoed by the recent work of *Chedin et al*., who demonstrated that issues of data quality persist even within prominent R-loop mapping studies (17). While these findings indicate the need for robust quality control methods in R-loop data analysis, no such techniques have yet been proposed to our knowledge.

In the present study, we profiled 693 publicly available R-loop mapping datasets. We devised methods to assess R-loop data quality control and applied these methods to identify high-quality R-loop mapping samples which allowed us to define consensus sites of R-loop formation, termed “R-loop regions” (RL regions). Meta-analysis of RL regions identified by different methods demonstrated stark contrasts between S9.6 and dRNH-based R-loop mapping results, supporting the proposed “Class I/II” distinction and revealing differing biological programs among the R-loops uncovered by each mapping approach. Moreover, we define constitutive and variable R-loop regions, finding that they differ with respect to pathways relevant for transcription and replication.

In this study, we also demonstrate novel approaches for the quality control and analysis of R-loop mapping data, which we provide in the collection of software tools, “RLSuite,” which accompany this work (see **Availability**).

## MATERIAL AND METHODS

### R-loop forming sequences (RLFS)

We predicted R-loop forming sequences using the *QmRLFS-finder* method (18) implemented as part of the *makeRLFSBeds.R* script in the accompanying data generation repository (see **Availability**). Briefly, we downloaded every genome for which gene annotations were available in “FASTA” format from the University of California Santa Cruz (UCSC) genome repository. Then we ran the QmRLFS-finder.py script to generate predictions. Finally, we converted the results table to “BED” format.

### Cataloguing R-loop mapping datasets

R-loop data was found by querying the Gene Expression Omnibus (GEO), Sequence Read Archive (SRA), ArrayExpress, and PubMed with the keywords “R-loop”, “R-loops”, “RNA:DNA hybrid”, “RNA:DNA hybrids”, “RNA-DNA hybrid”, “RNA-DNA hybrids”, “DRIP-Seq”, “R-ChIP”, “S9.6”, and “D210N”. We include all query results with R-loop mapping data publicly accessible via SRA. For every R-loop mapping sample, we recorded the following details:

1. The sample accession (GEO or SRA).
2. The accession for the corresponding genomic input sample (if available).
3. The sample condition (e.g., RNase H1 treated, “RNH”).
4. The mapping mode (e.g., “DRIP” for DNA-RNA immunoprecipitation sequencing (DRIP-Seq)).
5. The tissue of the sample. For cell lines, this was the cell line name (e.g., “HEK293”). For organs and whole organisms, the tissue name (e.g., “B-cell” for B-cells, “SC” for whole brewer’s yeast (Saccharomyces cerevisiae)).
6. The sample genotype, where relevant (e.g., “BRCA1” for samples derived from BRCA1 mutant breast cancer patients).
7. “Other,” a catchall for other relevant metadata provided by the original data submitters, such as drug treatment information.
8. The source publication PMID, if available.
9. The last author on the source publication, if available.

**Table S1** provides the full list of samples.

### Standardization of sample metadata

We standardized the manually curated catalogue via the custom scripts in the accompanying data generation repository (see **Availability**). First, the “condition” was binarized to one of the following labels: “POS” (samples which were expected to map R-loops, e.g., “D210N” in R-ChIP data) or “NEG” (samples which were not expected to map R-loops, e.g., “Input” for DRIP-Seq data). The mapping of “condition” to “label” relied on a manually curated regular expressions (REGEX) dictionary and simple string matching (**Table S1**). Additionally, we filtered the manifest to keep only R-loop mapping samples (except for bisulfite sequencing samples). Finally, we executed the RLPipes v0.9.0 software program to query the SRA database for missing metadata related to each sample (see **Availability**).

### R-loop mapping data standardization

To reprocess all available R-loop mapping datasets, we used a long-running computational pipeline that is available in its entirety with detailed instructions in the accompanying data generation repository (see **Availability**). For all upstream processing steps, we used RLPipes v0.9.0 (see **Availability**). We executed this pipeline on a remote server running Ubuntu 18.04 with 500GB of RAM, 192 cores, and a 10GiB/s internet connection available. The pipeline took one week to finish processing all samples. The primary steps of the pipeline involved all the following:

First, raw reads in SRA format were downloaded for each SRA run via the prefetch software from NCBI sra-tools. Then, reads were converted to “FASTQ” format using fastq-dump from sra-tools. Next, technical replicates were merged and interleaved (in the case of paired-end data) using reformat.sh from *bbtools* (19). Then, reads were trimmed and filtered with *fastp* (20), generating a quality report.

For R-loop mapping data, reads were aligned to the appropriate genome using bwa-mem2 (21). Then, alignments were filtered (minimum quality score of 10 enforced), sorted, and indexed using samtools (22) and duplicates were marked using samblaster (23). Then, peaks were called using macs3 (24) and coverage was calculated using deepTools (25).

The outputs of the pipeline were (A) peaks in “broadPeak” format, (B) coverage in “bigWig” format, (C) quality reports for alignments and raw reads, and (D) alignments in “BAM” format (see **Availability**).

### Normalization of read coverage

To calculate normalized coverage tracks for visualization purposes, we first calculated scaling factors using the trimmed mean of M-values (TMM) approach previously demonstrated for similar sequencing approaches (26) implemented via the edgeR R package (27). Then, we calculated the expected fragment sizes for each sample using the macs3 predictd method (24). We then calculated coverage using the bamCoverage command from deepTools (25) with arguments “--scaleFactor <scale_factor>-e <fragment_size> -ignoreDuplicates” where “<scale_factor>” is the scaling factor derived from edgeR and “<fragment_size>” is the predicted fragment size from macs3. The resulting tracks were visualized in the UCSC genome browser.

### R-loop mapping data quality control

Following data standardization, a quality control model was developed to filter out poor-quality samples prior to meta-analysis of R-loop consensus sites. This involved (1) R-loop forming sequences (RLFS) analysis, (2) quality model building, and (3) sample classification.

#### R-loop forming sequences analysis

R-loop forming sequences (RLFS) analysis was performed using a custom R script, rlfsAnalyze.R, available in the accompanying data generation repository (see **Availability**). The analysis script calls the analyzeRLFS function from the RLSeq R package (see **Availability**). This function implements a data-agnostic method for assessing the enrichment of R-loop mapping peaks within RLFS via the following procedure: Permutation testing is implemented via the permTest function from the regioneR (28) R package such that, for each permutation, R-loop peaks were randomized using the circularRandomizeRegions function and then the number of overlaps with RLFS was counted. 100 permutations are used by default to build an empirical null distribution for peak/RLFS overlap. Then the true number of overlaps from non-randomized peaks and RLFS was compared to the null distribution to calculate the Z-score and significance of enrichment. Finally, a Z-score distribution was calculated using the localZScore function from regioneR (28) within 5kb upstream and downstream of the meta-RLFS midpoint. For more detail, see the RLSeq reference (see **Availability**). For visualization purposes, the scaled Z score distributions were calculated for each peak set in the reprocessed data and visualized using locally estimated scatterplot smoothing (LOESS) regression with the standard error as the confidence interval.

#### Quality model building

The quality classification model was built using the models.R script in the accompanying data generation repository (see **Availability**). The process of model building is semi-supervised and utilizes a graphical user interface to allow operators with minimal coding experience to conveniently use it. The model building procedure involves the following steps: (A) The operator executes the models.R script from the command line, launching an interactive web interface that contains a table with SRA experiment ID and label (“POS” or “NEG”) for each sample along with a plot showing the Z-score distribution from the RLFS analysis of that sample. This interface also indicates samples that have been previously deliberated about. (B) The operator then decides which samples to exclude from the model building process. A sample should only be excluded when the Z-score distribution drastically differs from the expected Z-score distribution for its label (examples are provided to the operator in the web interface). (C) Once all samples with a label mismatch are selected, the operator clicks the “Build model” button which executes the model building process on the non-excluded data via the RMarkdown notebook, FFT-classifier.Rmd (available from the accompanying data generation repository). This notebook automatically executes the following steps: (C.1) The data are wrangled. (C.2) Then, the engineered features are calculated from the Z-score distributions (**Table 1**). The engineered features are then wrangled into matrix form with samples in rows and engineered features and label as the columns. The label in this case is “POS” (expected to map R-loops) or “NEG” (not expected to map R-loops). (C.3) Then, the feature matrix was standardized (centered and scaled) and transformed with the Yeo-Johnson normalization via the caret R package (29). (C.4) Then, the data was partitioned using a 50:25:25 (train:test:discovery) split. (C.5) The discovery set was analysed with the Boruta automated feature selection method via the Boruta R package (30) to select the parsimonious feature set used to train the classifier. (C.6) The training set was then used to train a stacked ensemble classifier from the caretEnsemble R package (31), with the following architecture:

- The ensemble meta model is a Random Forest classifier and the five base models in the stack are:

- Latent Dirichlet allocation
- Recursive partitioning
- Generalized linear model (logit)
- K-nearest neighbours
- Support vector machine (radial)
- 10-fold cross-validation repeated 5 times was implemented during training.

**Table 1.** Engineered features used in the quality model. Where “Z” is Z-score distribution, “ACF” is the autocorrelation function, and “FT” is the Fourier Transform (calculated via the fft function in R).

TABLE AND FIGURES LEGENDS
| ***Feature*** | ***Description*** |
| --- | --- |
| ***Z1*** | mean of Z |
| ***Z2*** | variance of Z |
| ***Zacf1*** | mean of ACF(Z) |
| ***Zacf2*** | variance of ACF(Z) |
| ***ReW1*** | mean of FT(Z) (real part) |
| ***ReW2*** | variance of FT(Z) (real part) |
| ***ImW1*** | mean of FT(Z) (imaginary part) |
| ***ImW2*** | variance of FT(Z) (imaginary part) |
| ***ReWacf1*** | mean of FT(ACF(Z)) (real part) |
| ***ReWacf2*** | variance of FT(ACF(Z)) (real part) |
| ***ImWacf1*** | mean of FT(ACF(Z)) (imaginary part) |
| ***ImWacf2*** | variance of FT(ACF(Z)) (imaginary part) |

(C.7) Finally, the model was evaluated on the test set with the *confusionMatrix* function from the caret R package (29).

#### Sample classification

Following model building, all samples were subsequently classified using the classifySamples.R script from the accompanying data generation repository (see **Availability**). This script primarily uses the predictCondition function from the RLSeq R package (see **Availability**). The procedure implements the following steps for each peak set in the reprocessed R-loop mapping data: (1) calculates the Fourier transform of the Z-score distribution, (2) reduces the dimensions to the engineered feature set (see Quality model building), (3) applies the pre-processing model to normalize these features (see Quality model building), and (4) implements the classifier to render a preliminary quality prediction (see Quality model building). Of note, the quality model alone does not provide the final prediction for each sample. Instead, the final prediction is “POS” (positive) only if all the following are true:

1. The RLFS Permutation test P value is significant (p < .05)
2. The Z-score distribution at 0bp is greater than 0 (see R-loop forming sequences analysis)
3. The Z-score distribution at 0bp is greater than the value at both −5000bp (start) and +5000bp (end) (see R-loop forming sequences analysis)
4. The quality model predicts a preliminary label of “POS.”

These conditions were chosen to ensure that a positive prediction from the model (based on the distribution of the Z score) also met baseline criteria which indicate enrichment in typical permutation testing (28). These criteria may prevent spurious predictions when the model is applied to data types on which it was not originally trained.

Finally, predictions were rendered for every sample for which peaks could be calculated (656/693 samples).

### QC model validation

To examine the internal and external validity of the predicted labels, several approaches were implemented: (A) RLFS consensus analysis, (B) genic feature enrichment analysis, and (C) sample correlation analysis. All 4 (prediction:label) conditions: POS:POS, POS:NEG, NEG:POS, NEG:NEG were compared in each approach.

#### Visualization of normalized coverage around MALAT1

This analysis involved four DRIP-Seq samples, “TCELL (Input)” (SRX2455193), “TCELL” (SRX2455189), “786-O (RNH)” (ERX3974965), and “786-O” (ERX3974964), in which both the “TCELL” samples were from the same study (SRP095885), and the “786-O” samples were also from the same study (ERP120322). Both “TCELL” samples and both “786-O” samples respectively were from the same study.

#### Feature enrichment analysis

Genic features were obtained via the TxDb.Hsapiens.UCSC.hg38.knownGene R package (32). The computationally predicted G4 quadruplex features were provided by Chariker et al. in their recent work (33). The experimentally determined G4 quadruplex features were determined by Chambers et al. via their previous chromatin immunoprecipitation (ChIP) sequencing study (GEO accession: GSE63874) (34). All samples for which peaks could be called (37 samples excluded) were subjected to feature enrichment analysis for genic features and G4 quadruplex features. This analysis applied the following procedure to test the enrichment of peaks within these features: Using the RLSeq R package (1) peaks were randomly down sampled to a maximum of 10,000 peaks, (2) down sampled peaks were overlapped with genic features, (3) then, intersect statistics were calculated using Fisher’s exact test via the valr R package (35), finally, (4) log2 odds ratios of each label:prediction group were compared using a Kruskal-Wallis test and pairwise Dunn pos-hoc tests with Bonferroni p value correction, all performed via the rstatix R package (36).

#### Correlation analysis

This analysis uses correlation of R-loop signal coverage around high-confidence R-loop sites to assess sample-sample similarity. To prepare the analysis, the coverage track (bigWig) files for each sample were quantified around high-confidence sites (as first described by Chedin et al. (17)) via the gsgCorr.R script in the accompanying data generation repository (see **Availability**). First, high-confidence R-loop sites were identified from ultra-long-read R-loop sequencing (SMRF-Seq) (37). Then, high-confidence sites were lifted from hg19 to hg38 and extended by 100kb bidirectionally. The sites were then binned into 1kb windows. Then, the coverage signal for every human R-loop mapping sample was summed over each bin to create a matrix of bins[i] x samples[j] containing the total signal of each sample within each bin [ij]. Finally, the cor function was used to calculate the Pearson correlation of the samples in the matrix. The results were visualized with the ComplexHeatmap R package (38).

### R-loop consensus analysis

This analysis identified human R-loop consensus sites within catalytically dead RNase H1 (dRNH) and S9.6-based samples. To identify general R-loop consensus regions (RL regions), the union of dRNH and S9.6 consensus sites was calculated. To obtain these data, the following procedure was implemented: First, samples were prepared for R-loop consensus analysis using the prepareConsensus.R script from the accompanying data generation repository (see **Availability**). The following criteria were used to select samples for R-loop consensus analysis: (1) label of “POS”, (2) prediction of “POS”, (3) at least 5000 peaks called under a p adjusted value of 0.05, (4) is a human sample. We chose to enforce a label of “POS” as “NEG”-labeled are subjected to treatments which should prevent robust R-loop mapping and may, therefore, introduce unwanted technical variance to the consensus analysis. We chose samples with at least 5000 peaks due to the need for random peak down sampling to 5000 ranges.

227 samples met the full criteria for inclusion in the analysis of the 336 “POS”-labeled human samples. For each included sample, peaks were randomly shuffled and down sampled to 5000 ranges to ensure high peak-count samples would not dominate the analysis. Finally, samples were partitioned based on whether they used dRNH or S9.6-based R-loop mapping. Then, each down-sampled peak set was processed further using the rlregions.smk workflow (built on snakemake) from the accompanying data generation repository (see **Availability**).

For S9.6 and dRNH samples, the following were performed separately: (1) The peak sets were intersected with 10bp genomic windows and the number of peak overlaps per window was counted (using bedtools (39)). (2) The resulting windows and counts were converted to a bedGraph file (and its binary bigWig file format) and sorted using bedtools (39). (3) macs3 bdgpeakcall (24) was then implemented with the options -g 1000 -c x where x=max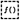(floor(.15*n_samples),5) and n_samples is the number of samples provided. These steps yielded (A) consensus coverage tracks (bigWig) and (B) consensus R-loop peaks (narrowPeak) for both S9.6 and dRNH samples.

Then, consensus RL Regions were generated using the peakUnion.R script from the accompanying data generation repository (see **Availability**). This script found the union of the peaks in the dRNH and S9.6 narrowPeak files, along with calculating the percent of samples within each group in which each peak appeared (conservation score) and averaging these scores in cases where dRNH and S9.6 peaks overlapped to create a peak union. Finally, RL regions and associated metadata were finalized using the finalizeRLRegions.R script from the accompanying data generation repository (see **Availability**). R-loops were filtered to remove any regions over 50kb in size, then every hg38 “POS”-labeled peak set was overlapped with the regions via the bed_intersect function from the valr R package (35) to yield a peak-RLRegion overlap matrix. This matrix was subsequently intersected with the RLBase sample metadata corresponding to each peak. Further details on additional metadata not used in this study are described in the RLHub R package reference (see **Availability**).

### Consensus site signal around genes

Consensus R-loop signal for dRNH and S9.6 was quantified around all hg38 ensemble genes using the computeMatrix function from the deepTools software program (25). The resulting data was scaled based on the number of samples in each group. The scaled signal was then scaled to a [0,1] range prior to plotting.

### Consensus site overlap in genes

Consensus sites for dRNH and S9.6 were length-standardized by finding the highest signal within the peaks and extending 250bp bidirectionally to form 500bp ranges. These peaks were then overlapped with transcript features. Finally, a priority rule was enforced such that peaks which overlapped two annotation types would be assigned to the one with the highest priority. The priority order was “TSS”, “TTS”, “fiveUTR”, “threeUTR”, “Exon”, “Intron”, “Intergenic”.

### R-loop region abundance calculation

R-loop region (RL region) abundance was calculated within each human sample in RLBase using the rlregionCountMat.R script from the accompanying data generation repository (see **Availability**).

RLBase sample alignment files (“BAM” format) were processed with featureCounts from the Rsubread R package to quantify the read counts from each within RL regions (40). Finally, the variance stabilizing transform (VST) was used to calculate normalized counts (41).

### Differential abundance of RL regions

Differential abundance between dRNH and S9.6 samples was calculated by the following procedure: (1) the count matrix was subset to contain only the RL regions found by both dRNH and S9.6 peaks, (2) the DESeq function from the DESeq2 R package (41) was applied to calculate differential abundance, (3) the enrichR R package (42) (and R interface to the enrichr web service (43)) was used to find the enrichment of gene ontology (GO) (44) terms within the genes overlapping differentially abundant RL regions.

Additionally, the prcomp function in R was used to calculate principal component analysis plots and the EnhancedVolcano R package (45) was used to generate the volcano plot. Finally, the pausing index for each gene (46) was calculated using the getPauseIndices function from the BRGenomics package (47). This calculation was performed using precision run-on (PRO) sequencing data in A549 cells downloaded from ENCODE (ENCFF275NOU and ENCFF979GYA) (48).

### Conservation analysis of RL regions

Conservation (percent of samples which detected an RL region) was ranked and bins were selected to partition RL regions by conservation percentiles. Binned RL regions were overlapped with genomic features and enrichment was calculated via the procedure described above (see Consensus site overlap in genes). Pathway enrichment was also performed via the procedure outline in the above section (Differential abundance of RL regions), with the only difference being that the ChEA (49), KEGG (50), and MSigDB (51) pathway databases were tested.

## RESULTS

### Curation of public R-loop mapping samples

The data in this study were hand-curated from public data repositories, yielding 693 R-loop mapping samples from 65 different studies (**Table S1**). The data are technologically and biologically diverse, representing 58 different tissue types, seven species, and 20 R-loop mapping modalities (**Table 2**, **Fig. S1A**). Moreover, the data includes both “POS” (expected to map R-loops) and “NEG” (not expected to map R-loops) labelled samples (**Table 2, Fig. S1A**). Examples of NEG-labelled samples include “WKKD” (RNase H1 that is incapable of both DNA binding and R-loop resolution (11)) R-ChIP data, RNase H1-treated DRIP-Seq, and genomic “Input” samples. Examples of POS-labelled samples include “D210N” (catalytically inactive RNase H1) R-ChIP data and typical S9.6 DRIP-Seq data. The scale and diversity of these data made them suitable as a basis for building our quality methods and for our subsequent meta-analyses.

**Table 2.** Characteristics of the R-loop mapping samples profiled in this study. For each, the number and percentage of samples are shown. Organism (UCSC ID): The species from which data were derived along with the UCSC genomic ID used in alignment. Mapping Mode: the type of R-loop mapping used (see **Table S1** for greater detail). IP Type: the type of immunoprecipitation (IP) used during mapping (based on the mode) where “dRNH” indicates catalytically dead RNase H1, “None” indicates no IP, “RNA-m6A” indicates methylated RNA, and “S9.6” indicates use of the S9.6 antibody. “Label”: the binarized form of the “condition” metadata assigned by the authors (see Methods); “POS” indicates the sample is expected to map R-loops and “NEG” indicates the sample is not expected to map R-loops.

|  | **Overall** |
| --- | --- |
| ***n*** | 693 |
| **Organism (UCSC ID) (%)** |  |
| *Roundworm (ce11)* | 6 (0.9) |
| *Zebrafish (danRer11)* | 7 (1.0) |
| *Fly (dm6)* | 30 (4.3) |
| *Chicken (galGal6)* | 8 (1.2) |
| *Human (hg38)* | 532 (76.8) |
| *Mouse (mm10)* | 77 (11.1) |
| *Yeast (sacCer3)* | 33 (4.8) |
| **Mapping Mode (%)** |  |
| *BisMapR* | 3 (0.4) |
| *DREAM* | 12 (1.7) |
| *DRIP* | 299 (43.1) |
| *DRIP-RNA-Seq* | 2 (0.3) |
| *DRIPc* | 40 (5.8) |
| *DRIPc-HBD* | 2 (0.3) |
| *DRIVE* | 1 (0.1) |
| *DRNA* | 10 (1.4) |
| *m6A-DIP* | 4 (0.6) |
| *MapR* | 54 (7.8) |
| *qDRIP* | 19 (2.7) |
| *R-ChIP* | 54 (7.8) |
| *RDIP* | 14 (2.0) |
| *RLCnT-HBD* | 13 (1.9) |
| *RLCnT-S96* | 16 (2.3) |
| *RNH-CnR* | 5 (0.7) |
| *RR-ChIP* | 5 (0.7) |
| *S1-DRIP* | 12 (1.7) |
| *sDRIP* | 49 (7.1) |
| *ssDRIP* | 79 (11.4) |
| **IP Type (%)** |  |
| *dRNH* | 137 (19.8) |
| *None* | 22 (3.2) |
| *RNA-m6A* | 4 (0.6) |
| *S9.6* | 530 (76.5) |
| **Label = POS (%)** | 440 (63.5) |

Following manual curation of R-loop mapping datasets, we developed a purpose-built R-loop pipeline tool called “RLPipes” and applied it to reprocess all datasets, yielding peaks, coverage, and other processed files (see **Availability**) (**Fig. 1A**).

**Figure 1.**
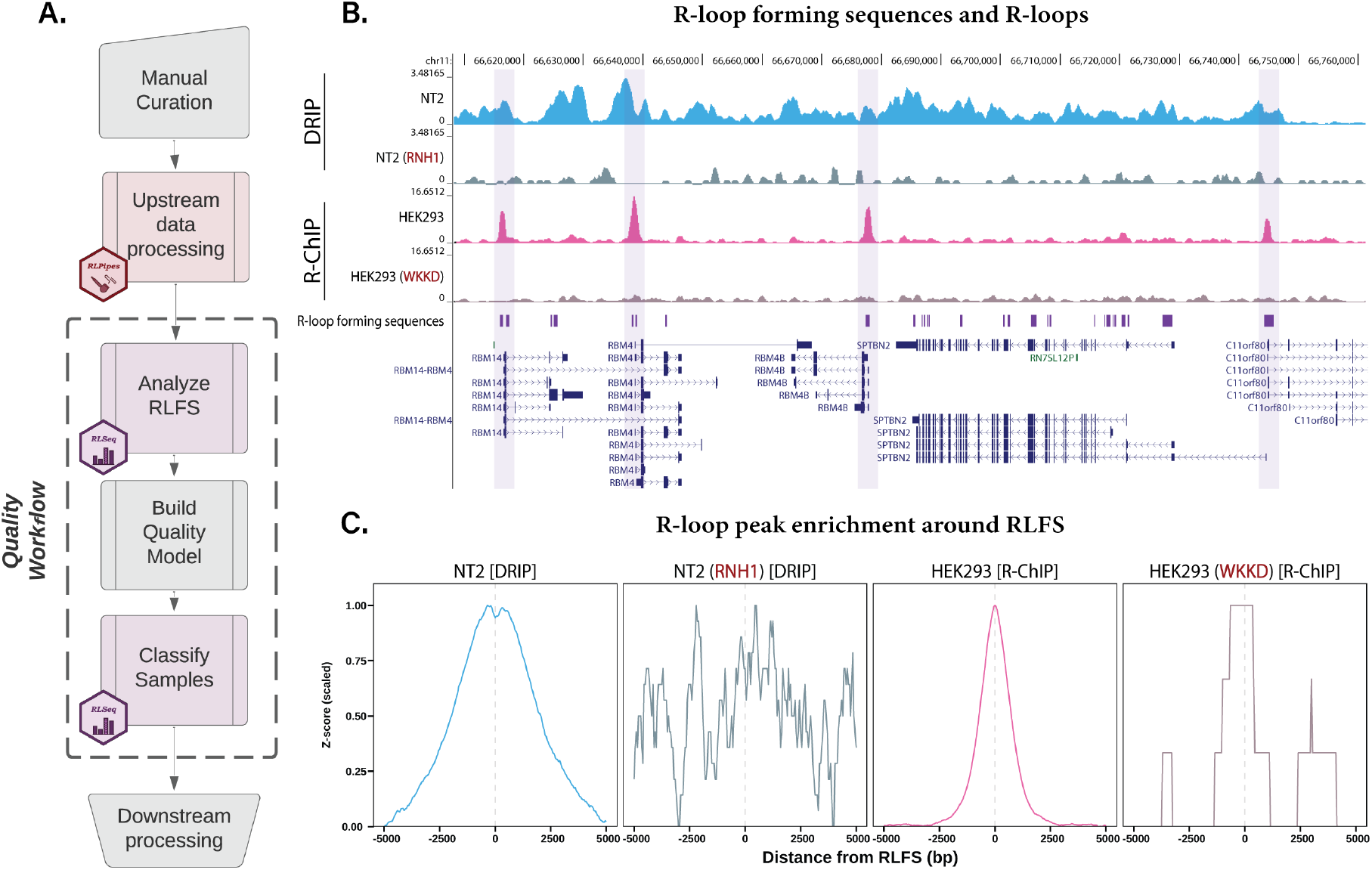
Reprocessing and standardization of R-loop mapping data. (A) The workflow used for reprocessed datasets along with icons indicating the software used at each step (see “**Availability**”). (B) A Genome Browser view of four representative samples (one “POS” and one “NEG” each from DRIP and R-ChIP modes) along with R-loop forming sequences (RLFS). Negative label samples are indicated with red text (e.g., “RNH1”). The genome browser session for this visualization is also provided within this manuscript (see **Availability**). The SRA IDs of these samples are SRX1025894 (“NT2”), SRX1025896 (“NT2 (RNH1)”), SRX2683605 (“HEK293”), SRX2675009 (“HEK293 (WKKD)”; WKKD: mutant RNase H1 without catalytic or DNA binding capabilities). (C) Metaplots of the four samples in (B) showing the enrichment of their called peaks around RLFS. The Y axis is the Z score after min-max scaling.

### The number of R-loops detected varies within and between modalities

Previous work has demonstrated discrepancies in the number of R-loop sites uncovered by different mapping methods (12). However, to our knowledge, no study to-date has quantified the extent of these differences. From our analysis of the numbers of peaks called from various techniques, we found discrepancies between modalities spanning multiple orders of magnitude (**Fig. S1B**). We found that DNA:RNA in vitro enrichment (DRIVE) sequencing produced the fewest peaks (337). However, only one DRIVE-Seq sample is publicly available at present (1). Following DRIVE-Seq, R-ChIP contains the fewest average peaks (~6 thousand). Strand-specific DRIP (ssDRIP) sequencing generated the greatest average number of peaks (~314 thousand). Moreover, we observed variability among the number of peaks even within mapping modalities (**Fig. S1B**). Taken together, these findings suggest that no single modality can confidently assess the locations of all R-loops within a sample. The results also reveal technical variance between samples belonging to the same mapping modality, suggesting potential issues with sample quality control across studies.

### R-loop forming sequences provide a suitable test of sample quality

Recent studies have indicated the presence of poor-quality R-loop mapping datasets within the literature (16, 17), but no tests of R-loop mapping accuracy have yet been proposed to our knowledge. To address this limitation, we developed a purpose-built quality control approach based on R-loop forming sequences (RLFS). RLFS are genomic regions which show favourability for R-loop formation (18, 52-55). We calculated RLFS across each genome and found that they agreed well with the R-loop mapping data we reprocessed (**Fig. 1B**). We then implemented permutation testing to assess the statistical enrichment of R-loop peaks within RLFS. From this analysis, we found a strong enrichment within RLFS for representative POS-labelled samples and a lack of enrichment within NEG-labelled samples (**Fig. 1C**). These findings indicated the suitability of using RLFS as a basis for assessing R-loop mapping data quality.

### Ensemble learning from RLFS provides accurate quality predictions

Evidence from our recent work and from Chedin et al. indicated that POS-labelled samples may sometimes fail to map R-loops as expected (16, 17). To address this concern, we performed RLFS analysis for every sample with peaks available (656 samples), yielding a p value and Z score distribution for each. As anticipated, we found some samples with a “POS” label, but which did not show expected enrichment around RLFS, based on Z score (**Fig. S2A**). This indicated the potential for using RLFS analysis to identify samples which fail to map R-loops as expected.

We reasoned that the p value from RLFS analysis would be sufficient for distinguishing sample quality. Unexpectedly, we found that, while many POS-labelled samples showed the expected Z-score distribution and significant p value, there were also examples of false positives in which the p value was significant, but poorly representative of both the assigned label and of the Z score distribution (**Fig. S2B**). This indicated the need for a more sophisticated test of sample quality from the RLFS analysis results, prompting our development of an ensemble learning model which could analyze the “noisiness” of the Z score distribution to provide additional discriminatory capability.

We built a classifier using a semi-supervised approach that involves a human operator deciding whether to exclude samples from the training set based on the alignment between the RLFS Z-score plot and the sample label (**Fig. S2C-D**). Of the 656 samples which had sufficient read depth for peak calling, the operator excluded 135 (20.6%) due to a mismatch between the sample label and the Z score distribution. Having selected the samples to discard, the operator triggered the automated model building script. Most features in the discovery set were automatically selected by Boruta for training (**Fig. S2E**). The ensemble learning model was then trained and evaluated internally via repeated cross validation, showing average performance above 0.9 specificity, 0.8 sensitivity, and 0.9 receiver operator characteristics area under the curve (ROC) for all base models (**Fig. S2F**). The final performance of the model was evaluated on a test set which the model had never previously seen. On this data, the model displayed an accuracy of 0.9043 (CI: 0.8353-0.9513; No Information Rate (NIR): 0.6435; P-Value [Acc > NIR]: 1.263e-10) (**Fig. S2G**), demonstrating the high accuracy of the model.

Finally, we combined the machine learning model with three other assessments to develop our final quality model. Briefly, a sample with a final prediction of “POS” must meet all the following criteria: (1) have a permutation testing p value below 0.05, (2) show a Z score above 0 at 0 bp in the distribution, (3) show a Z score at 0 bp which is greater than both the Z score at −5kb and +5kb, and (4) receive a prediction of “POS” from the machine learning model. We then applied the final “quality model” (criteria 1-4) to classify every reprocessed dataset in the present study (**Figure 2**).

**Figure 2:**
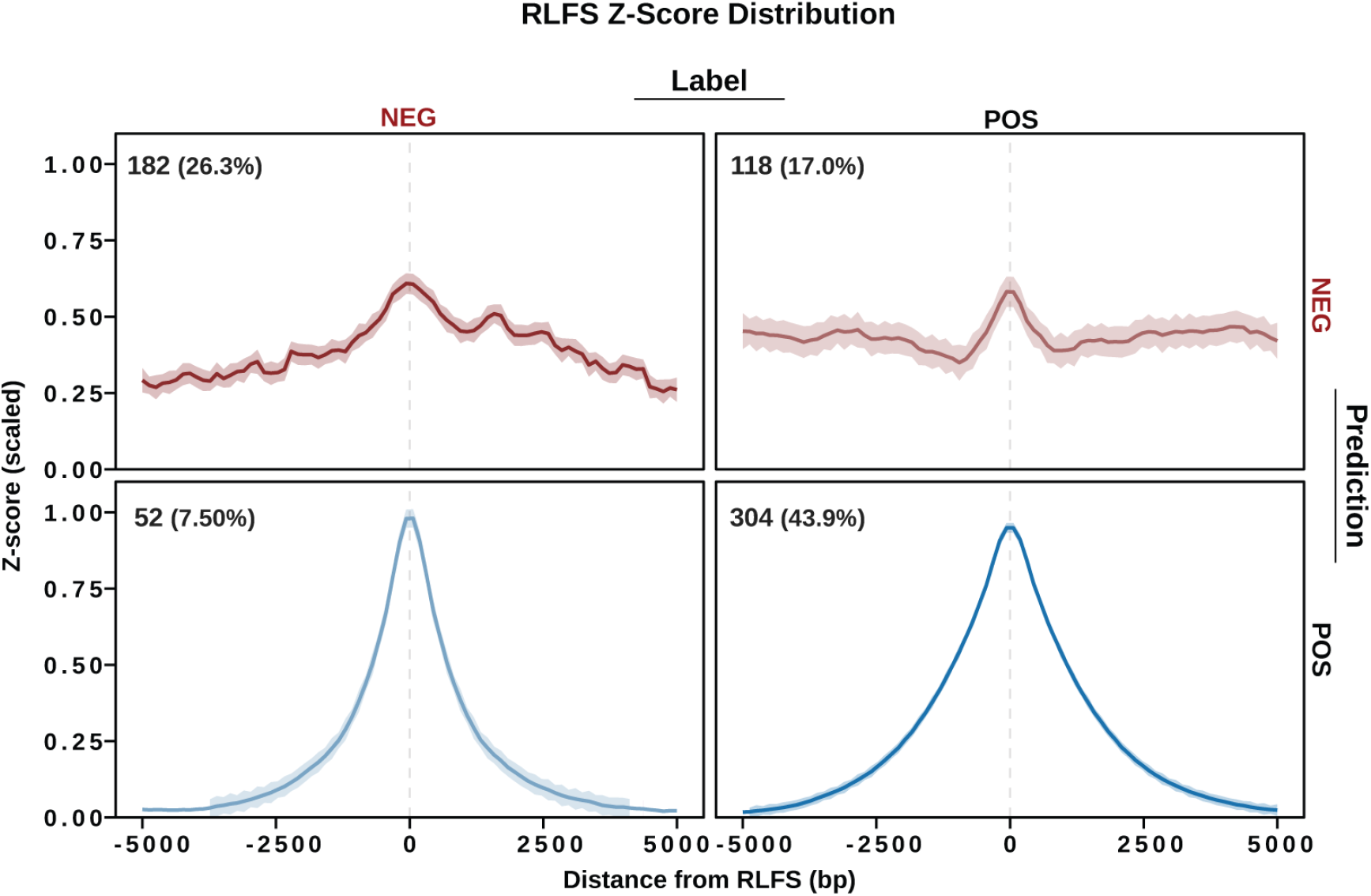
R-loop forming sequences analysis of prediction:label groups. Metaplots display the consensus signal among samples that were labeled “POS” (expected to map R-loops) or “NEG” (not expected to map R-loops) and samples which were predicted by the quality model to be “POS” or “NEG”. The plots show the LOESS regression line along with a confidence interval based on the standard error from LOESS regression. The annotations show the number and percentage of samples which are included.

### Validation of quality model reveals the prevalence of poor-quality R-loop mapping data

Unexpectedly, we found a discrepancy between the assigned label (based on author-supplied metadata) and the predicted label (based on our quality model) for many samples (**Figure 2**). Of the 234 NEG-labelled samples, the model classified 52 (22.22%) as “POS”, and, of the 422 POS-labelled samples, the model classified 118 (27.96%) “NEG”. To better understand this unexpected result, we divided the data by all prediction:label combinations (NEG:NEG, NEG:POS, POS:NEG, POS:POS) and calculated the consensus Z score distribution for each group. The results demonstrate that the classifications robustly represent the RLFS analysis of the data (**Figure 2, S3**). Coupled with the high accuracy of the ensemble model (**Fig. S2G**), these findings demonstrate that our quality model is *internally* valid.

We then proceeded to assess the *external* validity of our method through other quality approaches that did not involve RLFS analysis. The first approach was to visualize the normalized coverage tracks at sites of known R-loop formation. A representative depiction of four DRIP-Seq samples at a region near *MALAT1* (a non-coding RNA with constitutive R-loop formation (56)) shows that the model predictions distinguish between samples which display R-loop detection and those which do not (**Fig. 3A**). These results provided preliminary evidence for the external validity of our approach for determining whether a sample maps R-loops as expected.

**Figure 3.**
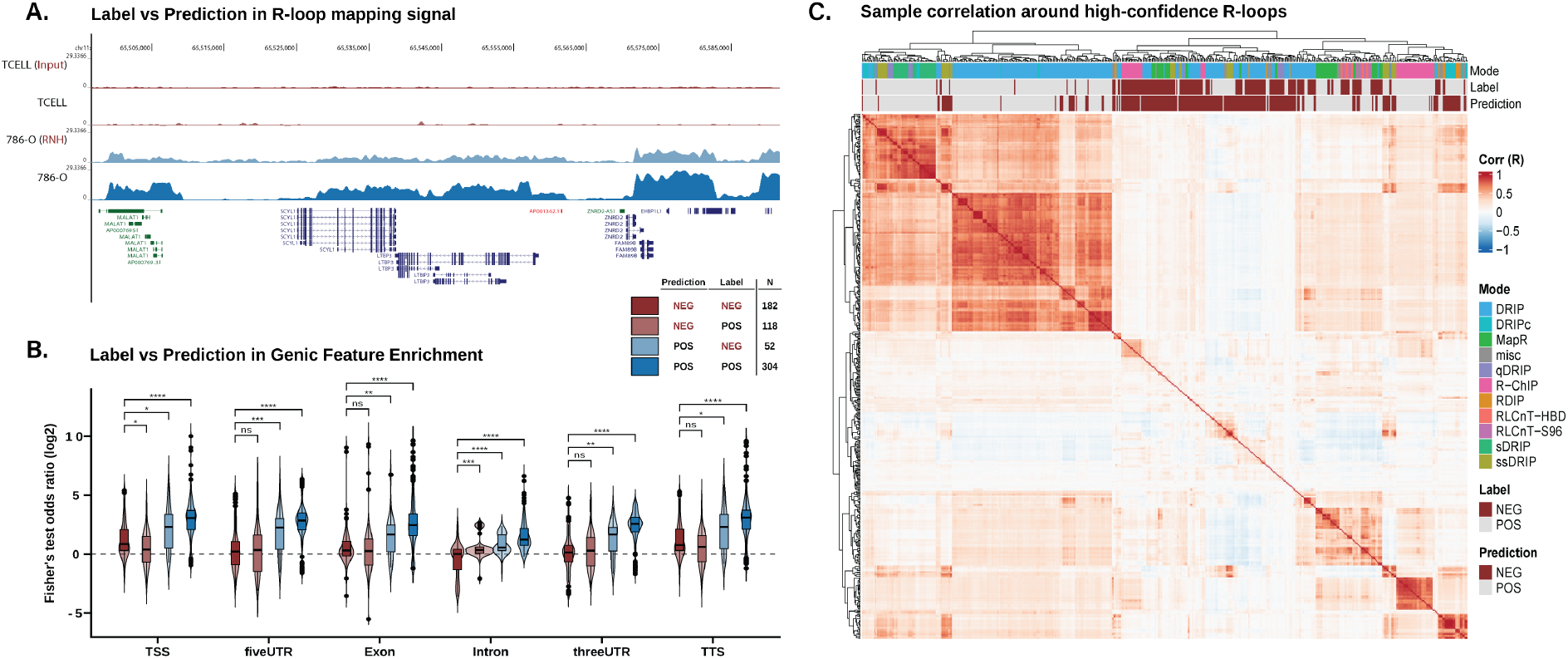
External validation of model predictions. (A) A Genome Browser image capture showing examples of DRIP-Sequencing datasets from two studies that exemplify all four possible model prediction versus sample label (prediction:label) combinations (“TCELL (Input)” [SRX2455193] - NEG:NEG; “TCELL” [SRX2455189] - NEG:POS; “786-O (RNH)” [ERX3974965] - POS:NEG; “786-O” [ERX3974964] - POS:POS). The genome browser session for this visualization is also provided within this manuscript (see **Availability**). (B) Violin/Box plots showing the distribution of Fisher’s exact test odds ratios within all reprocessed R-loop mapping samples, split by prediction:label combination and by the genomic feature on which testing was performed. Significance was determined via the Kruskal-Wallis test followed by Dunn post-hoc with Bonferroni correction. **** - p < .0001; *** - p < .001; ** - p < .01; * - p < .05; ns – p >= .05. (C) A heatmap showing the Pearson correlation (R) of reprocessed R-loop mapping samples around high-confidence R-loop sites. Dendrograms and row order show hierarchical clustering of samples and annotations show the prediction, label, and mode of each sample.

We then implemented enrichment testing for genic features, such as exons and introns, across all human samples in each prediction:label group. Given that R-loops are a by-product of normal transcription, successful R-loop mapping should result in an enrichment of peaks within genic features, as has been shown previously (11, 16, 57). From our analysis, we found that NEG-predicted samples, regardless of their original label, typically had low enrichment in all genic features (**Fig. 3B**). Moreover, we observed that the enrichment of genic features (with the exception of “Intron”) was not significantly greater in NEG:POS (prediction:label) samples compared to NEG:NEG samples (**Fig. 3B**). Conversely, POS:NEG showed a significant increase in enrichment in all features when compared to NEG:NEG (**Fig. 3B**). Together, these results support the external validity of our model predictions.

We then repeated the same analysis with both predicted and experimentally determined G4 quadruplexes (**Figure S4**). G4 quadruplexes are non-B DNA structures which can form on the displaced, non-template strand of R-loops and may promote R-loop stability (58–60). From feature enrichment analysis of G4 quadruplexes, we found that POS-predicted samples were significantly more enriched than NEG-predicted samples, regardless of prior label (**Figure S4**). These findings further demonstrate the external validity of the model predictions and their applicability to the analysis of R-loop mapping results in a biological context.

Finally, we implemented sample-sample correlation analysis to evaluate the agreement between the results obtained from the model prediction and the sample-level similarity as determined through correlation (**Fig. 3C**). As expected, we found that NEG-predicted samples tended to cluster together and display low correlation or no correlation with most POS-predicted samples (**Fig. 3C**). Likewise, we found that POS-predicted samples tended to cluster together, regardless of their original metadata label (**Fig. 3C**).

Taken together, these internal and external validations demonstrate that the method we present here is an accurate measure of R-loop mapping data quality. They also reveal the extent of poor R-loop data within published datasets, indicating that the approach described here will be of great benefit to future studies. Of note, we provide access to this quality method in both the RLSeq R package and the RLBase web application (see **Availability**).

### Consensus analysis reveals the locations and prevalence of R-loop formation

The locations and prevalence of R-loops genome-wide has been a subject of study since the first R-loop mapping modalities were described (1, 57). However, current estimates of where R-loops form and the percentage of the genome which they cover (3-13%) are derived from the analysis of small numbers of samples and mapping technologies (57, 61, 62). As we demonstrate here, there is variation in the number of peaks called both between and within modalities (**Fig. S1B**), indicating that the true prevalence and locations of R-loops remain unclear.

To address this gap in knowledge, we analysed high-confidence R-loop mapping samples to identify R-loop consensus regions (**Fig. 4A-D**), sites of robust R-loop formation across studies and modalities. Of note, our analysis workflow considers catalytically dead RNase H1 (dRNH) and S9.6 antibody (S9.6) R-loop mapping modalities independently prior to union analysis due to the hypothesized differences regarding the types of R-loops which these approaches map (13). Moreover, we provide the list of modalities belonging to the dRNH and S9.6 families within this manuscript (**Table S1**).

**Figure 4.**
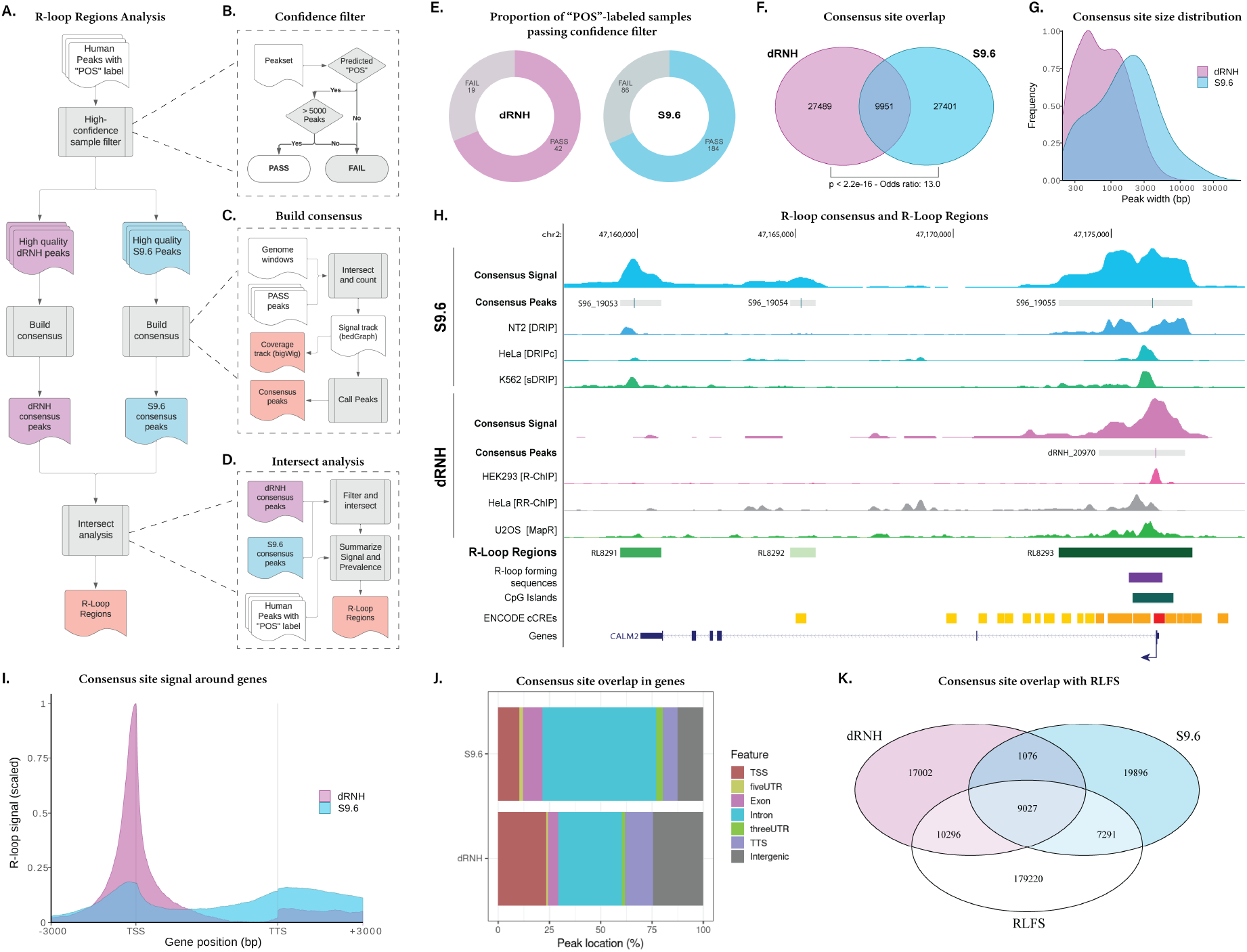
R-loop consensus analysis. (A) A flow diagram showing the workflow used for consensus analysis of R-loop sites in catalytically dead RNase H1 (dRNH) mapping samples and S9.6-based (S9.6) mapping samples. (B) A flow diagram showing the algorithm for selecting high-confidence samples to include in the analysis. (C) A flow diagram showing the workflow for building consensus R-loop signal and calling consensus peaks. (D) A flow diagram showing the workflow for intersect analysis, which yields R-Loop Regions. (E) Donut chart showing the proportion of “POS”-labeled dRNH and S9.6 samples passing the quality filters. (F) A Venn diagram showing the overlap of consensus sites derived from dRNH and S9.6-based mapping samples. P value and odds ratio from Fisher’s exact test. (G) Density plot showing the frequency of consensus peak width derived from S9.6 and dRNH-based mapping samples. X axis is log10 scaled. (H) Genome browser view of the strongest non-repetitive R-loop region (at the *CALM2* gene) with dRNH and S9.6 consensus signal, peaks, and signal tracks from representative S9.6 and dRNH R-loop mapping samples. The genome browser session for this visualization is also provided within this manuscript (see **Availability**). (I) Metaplot showing the R-loop consensus signal (scaled by total number of samples in dRNH and S9.6 respectively) around genes. (J) Annotation plot showing the proportion of summitted dRNH and S9.6 consensus sites within various genomic features. (K) Venn diagram comparing dRNH and S9.6 consensus sites to R-loop forming sequences (RLFS).

We first defined a high confidence set of samples based on the label, prediction, and number of peaks in the sample (**Fig. 4B**). We found that 42 of 61 human POS-labelled dRNH samples and 184 of 270 human POS-labelled S9.6 samples passed the quality filters (**Fig. 4E**). The analysis proceeded as described and yielded consensus sites and signal for dRNH and S9.6 separately, along with a consensus set of “R-loop Regions” (RL regions) from the union of S9.6 and dRNH consensus sites (see **Availability**). Of note, each valid “consensus site” was observed independently in at least 15% of all samples for both dRNH (6 out of 42 required) and S9.6 (27 out of 184 required). This constraint increases confidence that the consensus sites represent genuine R-loop formation.

We found that dRNH samples yielded 39,840 consensus sites while S9.6 samples yielded 37,449 consensus sites, together covering 4.32% of the human genome. Moreover, we calculated the overlap of these sites and found 9951 shared sites (**Fig. 4F**). This indicated that, despite having clear differences, dRNH was finding many of the same R-loops as S9.6. We then proceeded to evaluate the size distribution of the consensus sites (**Fig. 4G**). This analysis revealed that S9.6 peaks tend to be much larger, recapitulating previous reports (11). We then proceeded to examine these sites at *CALM2*, a gene containing one of the most mapped R-loops. Interestingly, this visualization showed that, while both S9.6 and dRNH peaks localize around the TSS, only signal from S9.6 was found in the gene body and around the TTS and PolyA regions (**Fig. 4H**). This finding led us to perform a meta-analysis of dRNH and S9.6 consensus sites and signal within genic features. From the results, we observed that while both S9.6 and dRNH show a preference for the TSS and TTS of genes, dRNH is more specific to these regions than S9.6 (**Fig. 4I**). Additionally, we calculated consensus site overlaps within genic features, demonstrating that a greater proportion of dRNH consensus sites occur in promoter regions and intergenic regions, while a greater proportion of S9.6 sites occur in Introns (**Fig. 4J**). From repeating this analysis with regions split into “Shared,” “dRNH-only,” and “S9.6-only” groups, we observed a co-localization of “Shared” consensus sites within promoter and TTS regions (**Figure S5**). Moreover, from this analysis, we observed an over-representation of S9.6-only consensus sites in intronic regions and of dRNH-sites in intergenic regions (**Figure S5**). Finally, we compared the location of R-loop consensus sites and RLFS, finding that most shared consensus sites co-localize with RLFS (**Fig. 4K**).

Taken together, these findings indicated that consensus R-loop sites recapitulate expected R-loop biology. Moreover, they reveal strong differences between dRNH and S9.6 mapping approaches with respect to localization in genic features as predicted by *Castillo-Guzman and Chedin* (13). Finally, this analysis produced a union set of R-loop regions which future studies can use for discrete analysis of R-loop mapping data (see **Availability**).

### dRNH- and S9.6-specific R-loop regions display key biological differences with respect to promoter pausing and ribosomal gene programs

In their recent work, *Castillo-Guzman and Chedin* predicted two previously unstudied classes of R-loops: “Class I” (R-loops resulting from promoter pausing) and “Class II” (R-loops resulting from transcriptional elongation) (13). They hypothesized that, while Class I R-loops are found by S9.6 and dRNH, Class II R-loops are more specific to S9.6 (13). In the above analysis of S9.6 and dRNH consensus sites, we found support for this hypothesis (**Fig. 4I-J**), but we did not yet address promoter pausing or the biological programs which are specific to Class I/II R-loops.

To address these questions, we analysed differential R-loop abundance for each high-confidence sample within the 9951 R-loop regions (RL regions) found in both S9.6 and dRNH consensus sites. We applied principal component analysis (PCA) to visualize the variance within this dataset (**Fig. 5A**). This revealed that dRNH and S9.6 samples diverge with respect to RL region abundance even though the sites analysed here represent sites found by both (**Fig. 5A**).

**Figure 5.**
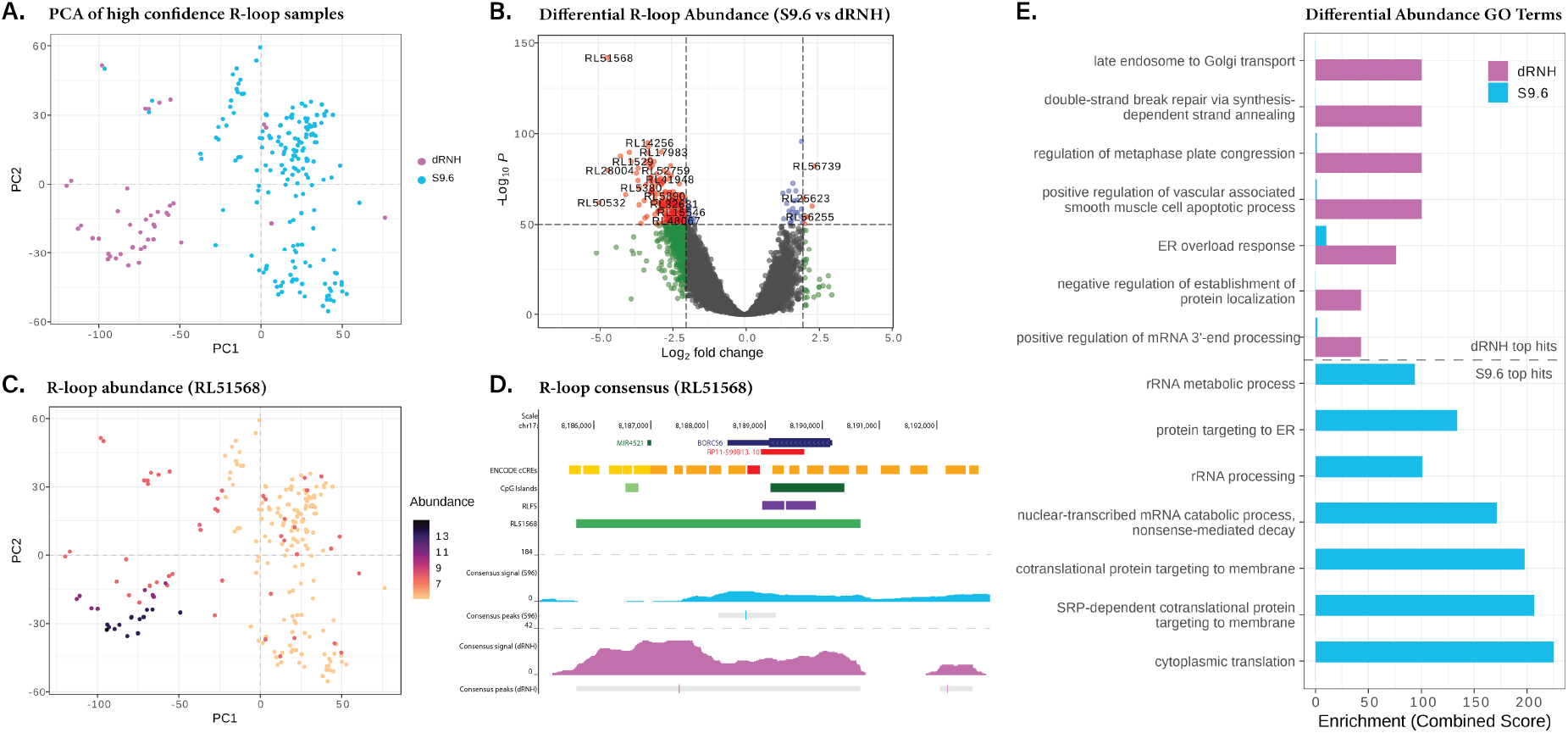
Differential abundance analysis of R-loop mapping signal. (A) A PCA plot showing the difference in R-loop profile between dRNH and S9.6-based approaches. (B) Volcano plot showing the top differentially abundant RL regions between dRNH and S9.6-based samples. (C) A PCA feature plot showing the top RL region (“RL51568”). (D) A genome browser view of S9.6 and dRNH consensus signal around RL51568. Coverage is scaled to the total number of high-confidence samples (S9.6: 184; dRNH: 42). (E) Top dRNH and S9.6-specific hits from GO term analysis of the genes associated with the top differentially abundant RL regions. Enrichment is measured by “Combined Score” such that higher indicates more strongly enriched.

Differential abundance analysis led to the identification of RL regions for which the S9.6-derived and dRNH-derived signal intensity were discordant (**Fig. 5B-C**) (see **Availability**). We visualized the top differentially abundant RL region (RL51568), demonstrating the difference within this region between S9.6 and dRNH methods (**Fig. 5D**).

To address the Class I/II hypothesis, we found the genes overlapping the dRNH-specific and S9.6-specific differential RL regions and calculated their pausing index (46). Of note, we subset these regions to only include those which overlap with the transcription start site (TSS) of genes. From the results, we observed that dRNH-specific genes show significantly greater pausing compared to S9.6-specific genes (2.13-fold) (**Fig. S6A**), supporting the validity of the Class I/II distinction.

To address the possibility that dRNH- and S9.6-specific R-loops occur in distinct biological programs, we found the genes overlapping the top one thousand in each and proceeded to implement GO term analysis. Unexpectedly, we found that, unlike dRNH, S9.6-based mapping methods robustly detected R-loops in genes related to ribosomal programs (**Fig. 5E**). To validate this finding, we examined the top 2000 dRNH and S9.6 consensus peaks and overlapped them with genes from the “KEGG Ribosome” pathway (as provided by the msigdbr R package (63)). Of 105 genes within the pathway, we found 58 in the top S9.6 consensus peaks and only 13 in dRNH consensus peaks (**Fig. S6B**). Moreover, in viewing the consensus signal around *RPL17* (a ribosomal gene and one of the top differential RL regions), we observed a strong difference between S9.6 and dRNH signal (**Fig. S6C**). These findings further indicate that S9.6 preferentially maps R-loops in many ribosomal genes compared to dRNH.

Taken together, these results highlight the validity of the proposed Class I/II distinction as dRNH seems to show a strong preference for genes with more promoter pausing. Moreover, they demonstrate biological differences between dRNH-based and S9.6-based R-loop mapping in several gene programs, especially those involving ribosomal genes.

### R-loop conservation analysis reveals biological differences between constitutive and variable R-loop regions

Finally, we elucidated the locations of “constitutive” RL regions (found consistently across samples) and of variable RL regions. To accomplish this, we calculated RL region conservation percentages, defined as the proportion of high-confidence R-loop mapping samples that identify each. From comparing dRNH and S9.6 conservation percentage distributions, we found that they tended to be similar, with a noticeable increase in conservation levels within sites found by both dRNH and S9.6 (**Fig. S7A**). We then proceeded to examine the distribution of conservation percentages across all RL regions and assigned them to bins for discrete analysis (**Fig. 6A**).

**Figure 6.**
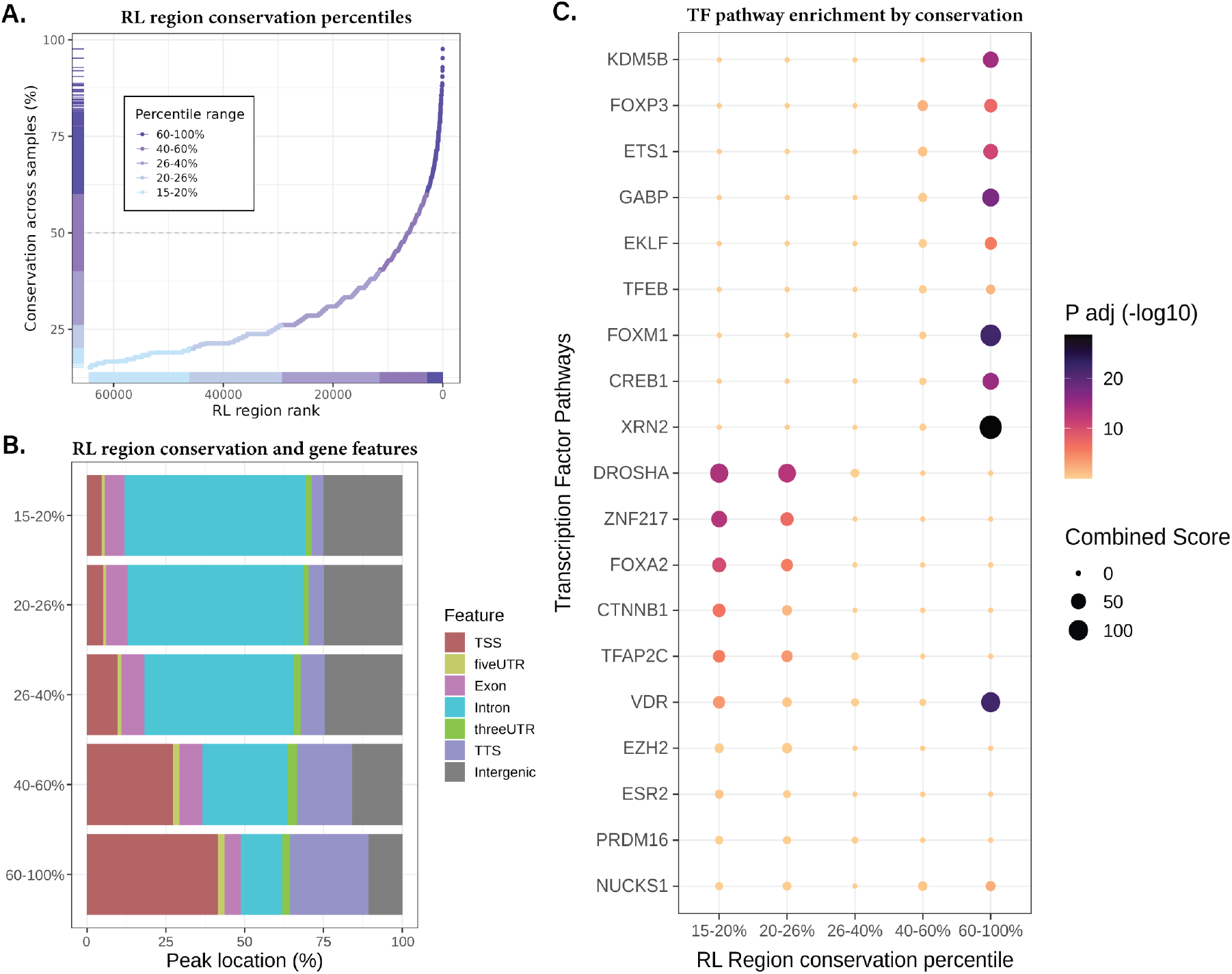
R-loop region conservation analysis. (A) Conservation rank plot showing all RL regions ordered by the percent of high-confidence samples in which they are found and binned by percentile ranges. (B) Annotation plot showing the gene features which overlap RL regions from various conservation percentile ranges. (C) Pathway enrichment plot showing the significance (via P adjusted value) and effect size (via Combined Score) of the “ChEA 2016” transcription factor pathway enrichment from the genes overlapping RL regions within each percentile.

Next, we examined the distribution of gene features overlapping the RL regions in each conservation bin. We found a strong association between high R-loop conservation and statistical enrichment within the transcription start site and transcription termination site of genes (**Fig. 6B**). Moreover, we saw a stronger enrichment of intron and intergenic features in variable (“15-20%”) RL regions (**Fig. 6B**). One explanation for this observation is that constitutive RL regions might associate with constitutively expressed “housekeeping genes” (64), while variable RL regions might overlap with more cell-type-specific genes. As expected, we observed a greater enrichment of housekeeping genes in more highly conserved bins (**Fig. S7B**). However, unexpectedly, the greatest enrichment was in the 40-60% conservation range and not the 60-100% range (**Fig. S7B**). This intriguing result led us to consider the possibility that conservation bins may represent biologically distinct R-loop programs that are partially independent from active gene expression.

To better understand the biological significance of constitutive and variable RL regions, we performed pathway enrichment on the genes overlapping each conservation bin. From an analysis of transcription factor binding pathways, we found stark divergence between constitutive and variable RL regions (**Fig. 6C**). The strongest hit from the constitutive regions was the “XRN2” target gene set. Notably, *XRN2* has been associated with R-loops in previous studies, particularly with respect to transcription termination (56). Conversely, the strongest hit from variable RL regions was the “DROSHA” target gene set. *DROSHA* is also a factor that has been previously studied in the context of R-loops (65). Moreover, we found *EZH2*, a core member of the polycomb repressive complex 2 (PRC2), was also associated with variable RL regions. Notably, PRC2 has been previously associated with R-loops in development-specific genes (66, 67). Curiously, we also observed the significant enrichment of the “VDR” target gene set in both constitutive and variable RL regions, but not in the bins between them (**Fig. 6C**). VDR, the vitamin D receptor, has not previously been associated with R-loop biology to our knowledge, further indicating the utility of this mode of analysis for revealing otherwise unknown associations within R-loop biology.

Having demonstrated the biological relevance of distinguishing between constitutive and variable RL regions, we proceeded to examine the KEGG and MSigDB pathways enriched within these groups. Curiously, we noticed the enrichment of “G2-M Checkpoint” in constitutive RL regions and “Mitotic Spindle” in variable RL regions (**Fig. S7C**), indicating the potential of different classes of R-loops to act in the regulation of cell division. We also observed the enrichment of “Spliceosome” and “RNA transport” in constitutive RL regions (**Fig. S7D**). Finally, we found enrichment of the “Adherens junction” pathway in the variable RL regions (**Fig. S7D**).

Taken together, these results demonstrate that R-loops span a range of conservation levels, with many being strongly constitutive and others being more variable across samples. They also reveal that constitutive and variable R-loops associate with divergent genomic features and gene programs, indicating they may represent biologically meaningful subclasses of R-loops which require further exploration.

## DISCUSSION

The role of R-loops in physiology and pathology remains a topic of much study at present (3). Outstanding questions concern the types of R-loops which exist, their prevalence, their degree of conservation, their associations with epigenomic factors and processes, and the differences between various R-loop mapping approaches. Many of these questions cannot be faithfully answered by a single dataset, but instead must be gleaned through meta-analysis of hundreds of samples representing diverse biological and technical conditions. In the present study, we reprocessed and standardized 693 publicly available R-loop mapping datasets, developing the largest such resource to date. From these data, we developed a new and highly accurate method of quality control and implemented it to identify high-confidence R-loop mapping samples suitable for meta-analysis. From this analysis, we defined R-loop consensus sites and elucidated the differences between S9.6 and catalytically dead RNase H1 (dRNH) mapping approaches. Finally, we derived a set of “R-loop regions” (RL regions) from the union of dRNH and S9.6 samples and performed a conservation analysis which revealed the biological significance of constitutive and variable R-loops. The following sections summarize the key findings and provide additional context for their relevance to the field.

### A robust quality model reveals the prevalence of poor-quality data in the literature

Within the R-loop mapping field, no methods to assess whether an individual sample has robustly mapped R-loops currently exists. Instead, studies rely only upon quality methods which are generic to all sequencing datasets, such as FastQC (11, 68). Previous meta-analyses of R-loop mapping datasets relied upon comparisons with data from the literature to indirectly gauge quality (16, 17), an approach which could be confounded by systematic biases affecting large numbers of studies. From these limitations, and the previous evidence of inconsistent R-loop data quality (16, 17), we identified a clear need for a new R-loop mapping QC method that could benefit future researchers while also revealing the prevalence of high- and low-quality R-loop data within the field.

R-loop forming sequences (RLFS) are genomic regions which show favourability for R-loop formation (18, 52-55). They are computationally predicted from genomic sequence alone and show consistency with R-loop mapping data and with other computational prediction approaches, such as skewr (G or C skew prediction) (1, 2, 18). We calculated RLFS across each genome and found that they agreed well with the R-loop mapping data we reprocessed (**Figure 1**), indicating their utility as the basis for our quality control methodology. We then developed a quality model based on ensemble learning which uses the outputs from RLFS analysis to classify samples as “POS” (robust R-loop mapping) or “NEG” (poor R-loop mapping). This approach demonstrated excellent accuracy (**Fig. S2E-F**), along with both internal and external validity (**Figure 2, 3, S4**). Unexpectedly, we found many samples (24.5%) with mismatches between the expected label (from sample metadata) and the predicted label from our model (**Figure 2**). Taken together, these results indicate the troubling extent of poor R-loop data within published datasets, indicating that the approach described here will be of great benefit to future studies. Of note, we provide access to the method described here via the RLSeq R package and the RLBase web application (see **Availability**).

### R-loop consensus site analysis reveals the sites of robust R-loop formation

While it has been proposed that roughly 3-13% of the genome may contain R-loops, these findings were derived from an analysis of peaks from a limited number of S9.6-based R-loop mapping samples (57, 61, 62). From our analysis, we found stark discrepancies in the number of peaks called between studies and between R-loop mapping modalities (**Fig. S1B**). These findings indicate that efforts to faithfully assess R-loop locations and prevalence require a multi-study and multi-modality consensus analysis. From our meta-analysis, we identified robust sites of consensus R-loop formation which cover 4.32% of the human genome, termed “R-loop regions” (RL regions). These regions reveal the locations of R-loop formation and provide a discrete set of genomic annotations for future studies to use in the analysis of their R-loop mapping data.

### dRNH and S9.6 differentially map Class I/II R-loops in distinctive biological programs

*Castillo-Guzman and Chedin* recently proposed a “Class I/II” distinction where “Class I” refers to R-loops which result from promoter-proximal pausing, and “Class II” refers to R-loops which result from transcriptional elongation (13). They hypothesize that dRNH may preferentially find “Class I” R-loops whereas S9.6 may find both “Class I” and “Class II” R-loops (13). However, no bioinformatics analysis has yet demonstrated the existence of these R-loop classes genome wide.

From our analysis of dRNH and S9.6 consensus sites (**Figure 4, 5**), we found that S9.6-based R-loop mapping tended to produce larger peaks across the gene body while dRNH peaks tended to be smaller and more localized to promoter, TSS, or TTS sites as predicted by the Class I/II distinction. Our subsequent analysis showed that dRNH-specific R-loop regions occur in genes which have a significantly higher pausing index (**Fig. S6A**). Altogether, these findings support the Class I/II hypothesis.

Given that S9.6 and dRNH appeared to map biologically distinct R-loops, we also decided to analyze the differences in gene programs found by each (**Fig. 5E**). Most striking was the observation that R-loop consensus sites found by S9.6, but not by dRNH, were strongly associated with pathways relevant for ribosome biogenesis and translation (**Fig. 5E, 6B-C**). At present, it is unclear why S9.6 detects R-loops in ribosomal genes more efficiently or how this relates to the differences between Class I and Class II R-loops, indicating the need for further analysis by future studies.

### Constitutive and variable R-loops occur in distinct biological pathways

Another outstanding question in the R-loop field relates to the dynamics of constitutive R-loops (those which are consistently abundant across biological conditions) and variable R-loops (those which form under specific biological conditions). To address this question, we analysed constitutive and variable sites of R-loop formation (**Figure 6, S7**). From our analysis, we found a strong relationship between RL region conservation and enrichment within TSS and TTS regions of genes. Because R-loops tend to form as a result of transcription in some genes, we decided to test whether high R-loop conservation was associated with “housekeeping genes”, genes which are thought to be constitutively expressed in most cell types (64). Curiously, this analysis revealed that, while constitutive R-loops co-localize with many housekeeping genes, these genes were not as strongly associated with the most highly conserved RL regions (**Fig. S7B**). This unexpected result may indicate that, while high R-loop conservation is associated with constitutive gene expression, there may be constitutive R-loops which form more independently from gene expression. Additionally, from pathway enrichment we revealed a striking divergence in the biological programs and pathways enriched within constitutive and variable RL regions (**Fig. 6C, S7 C-D**). These findings indicate new and interesting directions for future R-loop studies to pursue.

### Limitations

#### Quality model training data

While the quality control method proposed in this study is both internally and externally valid (**Figure 2, 3, S2, S3, S4**), it may not be well-suited to assess the R-loop mapping techniques introduced in the future or those for which limited data is currently available. As of this writing, there are 23 different techniques for R-loop mapping, 14 of which were introduced since 2019, with seven introduced in 2021 alone (**Table S1**). Many of these techniques rely upon approaches that do not use either S9.6 or dRNH, such as m6A-DIP (69) and could pose unforeseen challenges for our quality model. Moreover, several of these techniques have few samples from a small number of studies, making it difficult to generalize about them. Finally, some species have few R-loop mapping samples available. To address these limitations, we have developed an online learning scheme whereby an operator can, with minimal training and software experience, rebuild the model in a matter of minutes to include new datasets. To facilitate this process, we developed a user-friendly web-browser interface for interacting with the model-building software (**Fig. S2A-D**). This workflow offers a convenient and automatic model building scheme that will make it easy for us to add new data as it becomes available so that our model continually improves over time.

#### Consensus analysis

We derived R-loop consensus sites from 42 dRNH and 184 S9.6 high-confidence R-loop mapping samples. This represents all the available human data which passed our quality thresholds. However, it still indicates a significant imbalance between dRNH and S9.6, leading to potential inconsistencies in the accuracy of our findings. As we explain above, we have made provisions to ensure that the process of incorporating new data is as convenient and automated as possible. We expect this will make it easy for us to regularly update the database and, along with it, the robustness of the biological interpretation of these data.

Another caveat for the joint analysis of S9.6 and dRNH-derived data is that there are large discrepancies between the size of peaks, coverage regions, and consensus sites between them (**Fig. 1B-C, 4G**). S9.6-derived peaks, coverage, and consensus sites tend to be much wider than those from dRNH (**Fig. 1B-C, 4G**). This may be due to the prevalence of DRIP-Seq data in the S9.6 group. Because DRIP-Seq relies upon restriction enzyme digestion, it may be biased toward larger peaks. Analyses for which this poses an immediate problem are those involving enrichment in genomic features when a feature priority is used (Fig. 4J, 6B) as larger peaks may overlap with more genomic features, leading to spurious over-enrichment in top-priority features. Additionally, for analyses which involve finding genes that overlap with R-loop sites, larger peaks may lead to overlaps with many more genes than shorter peaks would find. To address this issue, we forced the peaks to a uniform width centered around peak “summits” (the areas of highest enrichment from peak calling) for these analyses.

#### Bisulfite samples and stranded data analysis

We excluded bisulfite samples from our analysis due to the added complexity of processing them and the small number of available samples (only 27 in total). However, these techniques may be useful in providing nucleotide-level R-loop mapping. Therefore, in future versions of the software used for this analysis, we will include methods for analysing these data types. Another limitation of this analysis is that it does not incorporate strand assignment for any of the strand-specific sequencing techniques included in the dataset. This is intentional as DRIP-Sequencing (an *unstranded* mapping method) is still the most well-represented modality and all analyses require conversion to unstranded data for compatibility with it. However, as stranded techniques gain in popularity and begin to outpace DRIP-Seq, we anticipate that future versions of the software used for processing these data will incorporate strand information. Therefore, we expect to eventually update the analyses presented in this study with stranded data to reveal higher-resolution insights into R-loop dynamics.

## CONCLUSION

In this study, we reprocessed and standardized 693 public R-loop mapping samples, introducing a novel method of quality control to assess them. From the results, we demonstrate the widespread inconsistency of dataset quality within the literature and highlight the need for future studies to adopt our quality control approach. Moreover, from the high-confidence R-loop samples identified herein, we proceeded to define RL regions, sites of R-loop consensus across studies. From an analysis of these regions, we revealed the stark differences between the R-loops found by dRNH and S9.6-based techniques. Furthermore, we uncovered new biologically relevant subtypes of R-loops based on their conservation across samples. Taken together, this study introduces new methods and exciting future research questions for the R-loop mapping field.

## AVAILABILITY

### Data availability

No new raw data were generated as part of this study. All processed data files generated as part of this study are provided via RLHub and RLBase: https://gccri.bishop-lab.uthscsa.edu/rlsuite/. The list of R-loop regions (RL regions) and the list of differential RL regions between S9.6 and dRNH samples are provided on Figshare: https://doi.org/10.6084/m9.figshare.16920475.v1. UCSC genome browser sessions shown in this study are also provided:

1. Fig. 1B: https://genome.ucsc.edu/s/millerh1%40livemail.uthscsa.edu/Fig_1B_rloops_rlfs
2. Fig. 3A: https://genome.ucsc.edu/s/millerh1%40livemail.uthscsa.edu/Fig_3A_Malat1
3. Fig. 4H: https://genome.ucsc.edu/s/millerh1%40livemail.uthscsa.edu/Fig_4H_rl_consensus
4. Fig. 5D: https://genome.ucsc.edu/s/millerh1%40livemail.uthscsa.edu/Fig_5D_RL51568
5. Fig. S6B: https://genome.ucsc.edu/s/millerh1%40livemail.uthscsa.edu/Fig_S6C_ribogenes

### Code availability

The code used in this study is provided publicly on GitHub. The repository containing the workflow for recreating all upstream data analyses is provided here: https://github.com/Bishop-Laboratory/RLBase-data. The repository containing the code used to generate all figures in this manuscript is provided here: https://github.com/Bishop-Laboratory/RLoop-methods-and-analysis-2021. The software developed as part of the present study may be accessed here: https://gccri.bishop-lab.uthscsa.edu/rlsuite/. An interactive web application for exploring the data resource developed herein and analysing user-supplied R-loop mapping peaks (RLBase) is available here: https://gccri.bishop-lab.uthscsa.edu/rlbase/.

## ACCESSION NUMBERS

All accessions for data used in this study are listed in **Table S1**.

## SUPPLEMENTARY DATA

Supplementary Data are available at NAR online.

## ACKNOWLEDGEMENT

We want to thank all those who contributed to the development of the software used during this study, particularly the contributors to “RLSuite.” We want to thank Lori Kern at Bioconductor for her review of RLSeq and RLHub. We want to also thank Simon Bray at Bioconda for his review of RLPipes.

