## Supplemental Figures for "Quality-controlled R-loop meta-analysis reveals the characteristics of R-Loop consensus regions"

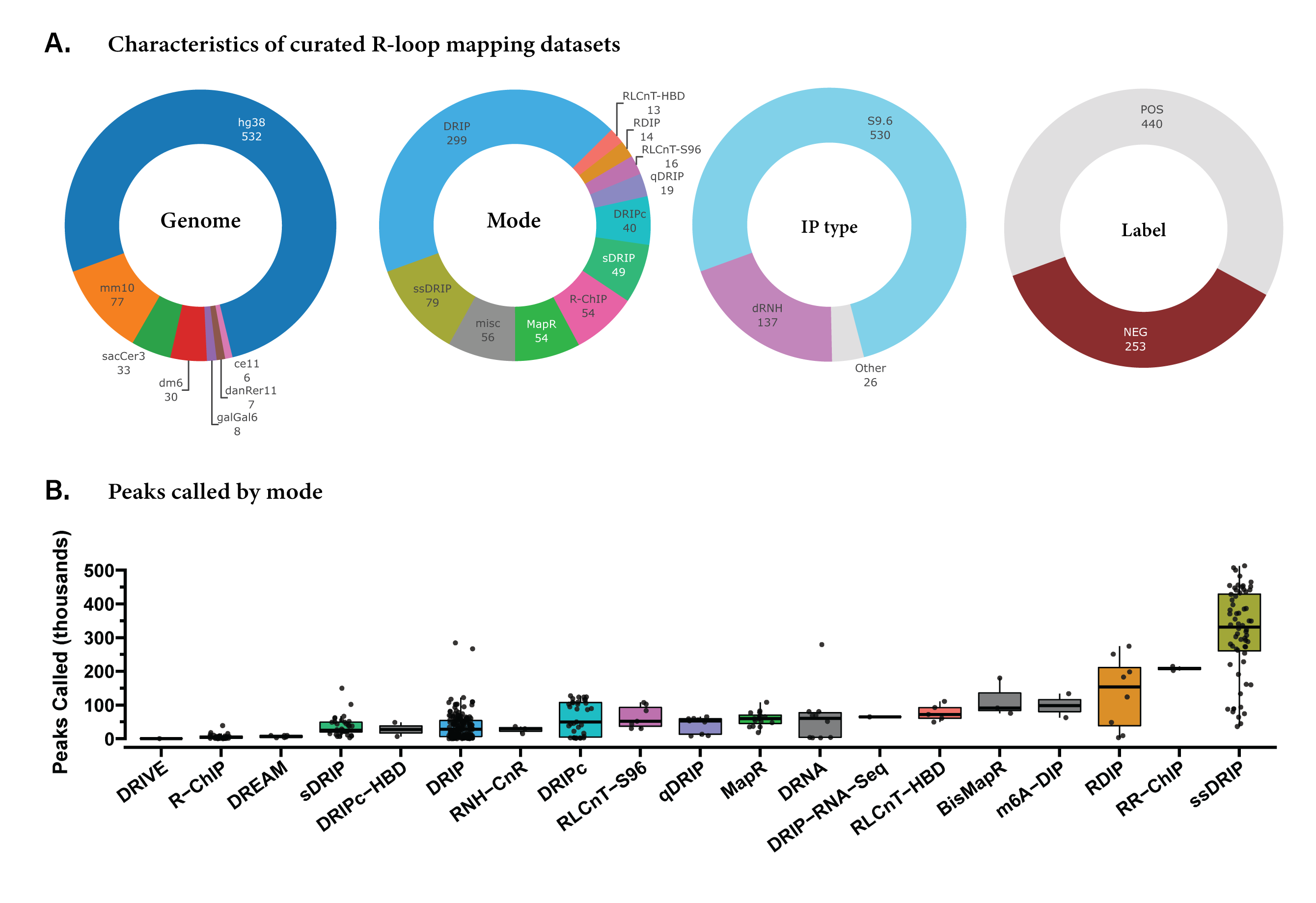

**Figure S1. Overview of reprocessed samples.** (A) Donut charts summarizing the proportion of different genomes represented in the dataset (“Genome”), R-loop mapping modes in the dataset (“Mode”), the proportion of immunoprecipitation types for those mores (“IP type”), and the binarized labels which correspond to the sample metadata and indicate whether a sample is expected to map R-loops (“POS”) or is not expected to map R-loops (“NEG”) (“Label”). (B) Box+jitter plots showing the number of peaks called in the reprocessed samples, split by mapping mode. Y axis shows the number of peaks in thousands.

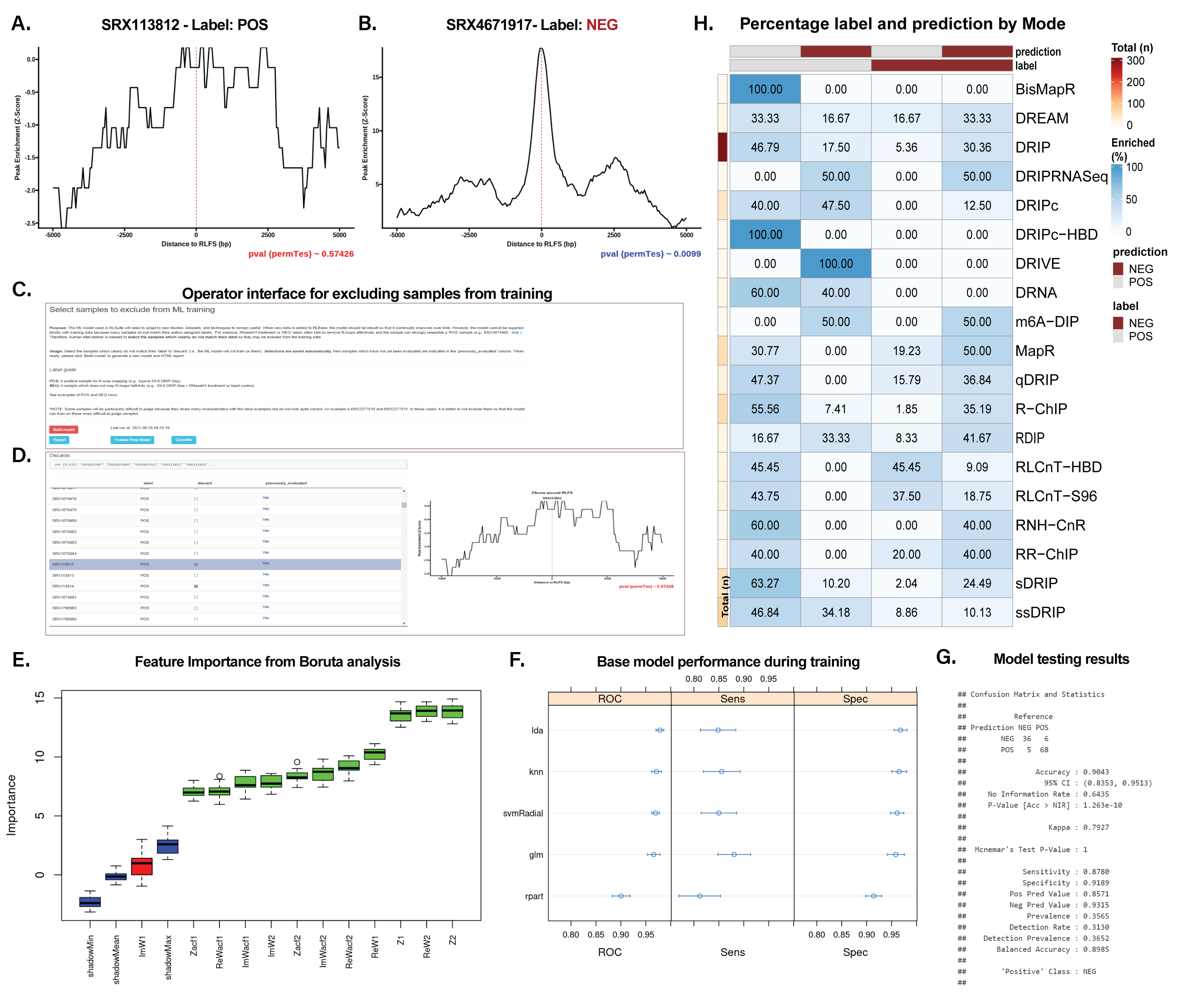

**Figure S2. Model training and results.** (A) Discordant RLFS analysis result in which Z score distribution does not match the “POS” label for this sample. (B) Discordant RLFS analysis results in which the permutation testing P value (p < 0.0099) does not match the “NEG” label and it does not appropriately represent the Z score distribution. (C-D) The operator interface for semi-supervised model training. (C) Instructions and links to examples for the operator to use. The “Build Model” button will automatically launch the model building script and upload the results. This section also contains the timestamp from the last model building run. (D) A data table with one entry per sample that provides the sample label, a checkbox which is used to add the sample to the discard, and a column “previously_evaluated” which indicates whether the sample was previously seen by an operator before the last model version was built. The row selected in this table also controls which plot is shown. The plot is an RLFS Z-score distribution plot corresponding to the sample selected in the table. This plot shows a sample which was labeled “POS” (expected to map R-loops), but which an operator chose to discard as the Z score distribution did not fit the assigned label. The p value annotation relates to the p value from permutation testing (see Methods). (E) The box plot showing the feature importance of each engineered feature in the discovery set as determined by analysis with Boruta (see Methods). (F) A model evaluation plot showing the receiver operator characteristics (ROC), the sensitivity (Sens), and the specificity (Spec) for each base model in the classifier during training. Confidence intervals were obtained by repeated cross-validation (see Methods) and indicate the range of potential values for each model. (G) The model performance on the test set as directly reported by the model. This includes a confusion matrix which shows the number of correct and incorrect predictions. (H) A heatmap showing the proportion of prediction:label combinations for all samples by mode.

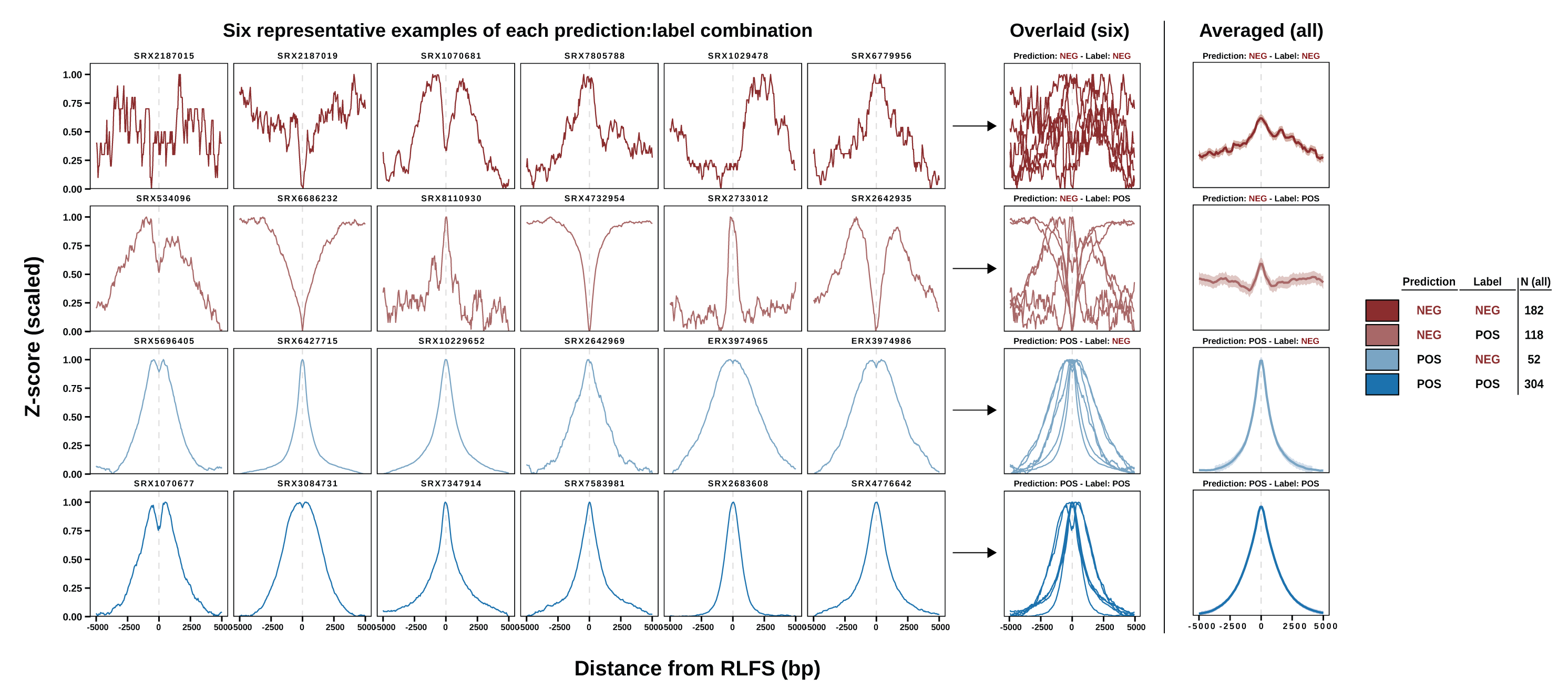

**Figure S3. Examples of each prediction:label combination and aggregation.** The grid of figures labeled “Six representative examples of each prediction:label combination” display RLFS metaplots which show the enrichment of the peaks within each sample around RLFS. The column of figures labeled “Overlaid (six)” shows the overlay of the six figures in each show. The column labeled “Averaged (all)” shows the summarization of all data within each prediction:label group (see legend for n), calculated via loess regression.

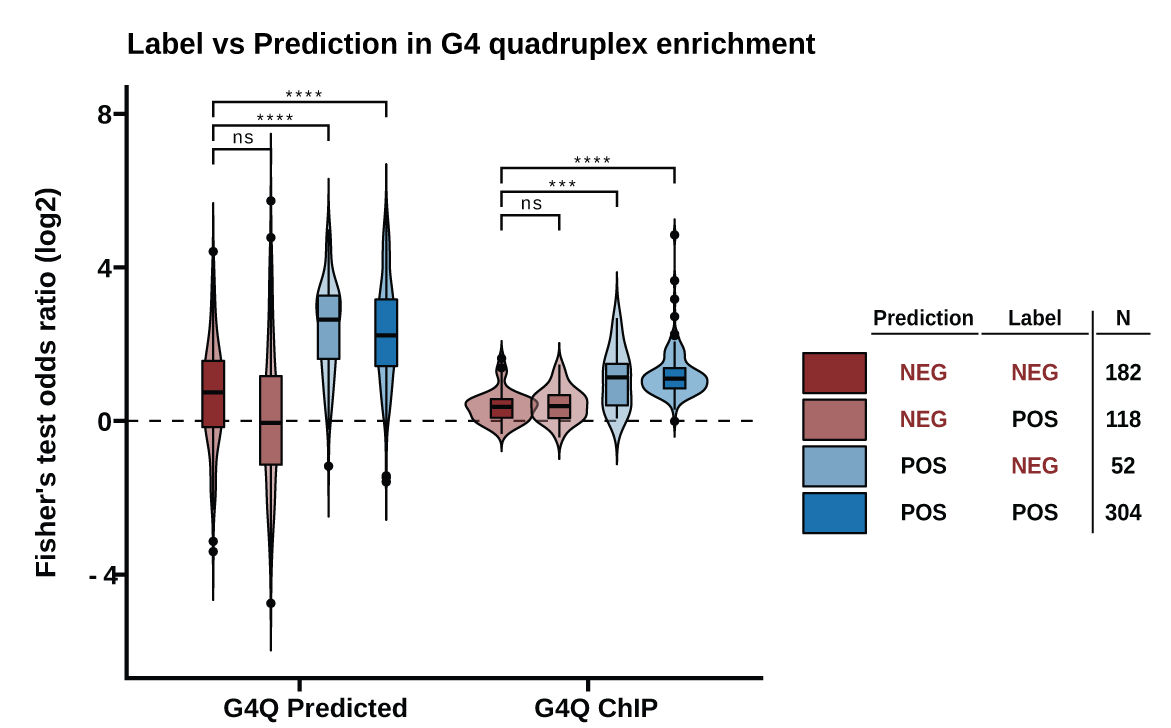

**Figure S4. Enrichment test in G4 quadruplex features.** Violin/Box plots showing the distribution of Fisher’s exact test odds ratios within all reprocessed R-loop mapping samples, split by prediction:label combination, for enrichment testing within predicted and experimentally-determined G4 quadruplex sites. Significance was determined via the Kruskal-Wallis test followed by Dunn post-hoc with Bonferroni correction. The legend shows the number of samples within each prediction:label combination. **** - p < .0001; *** - p < .001; ** - p < .01; * - p < .05; ns – p >= .05.

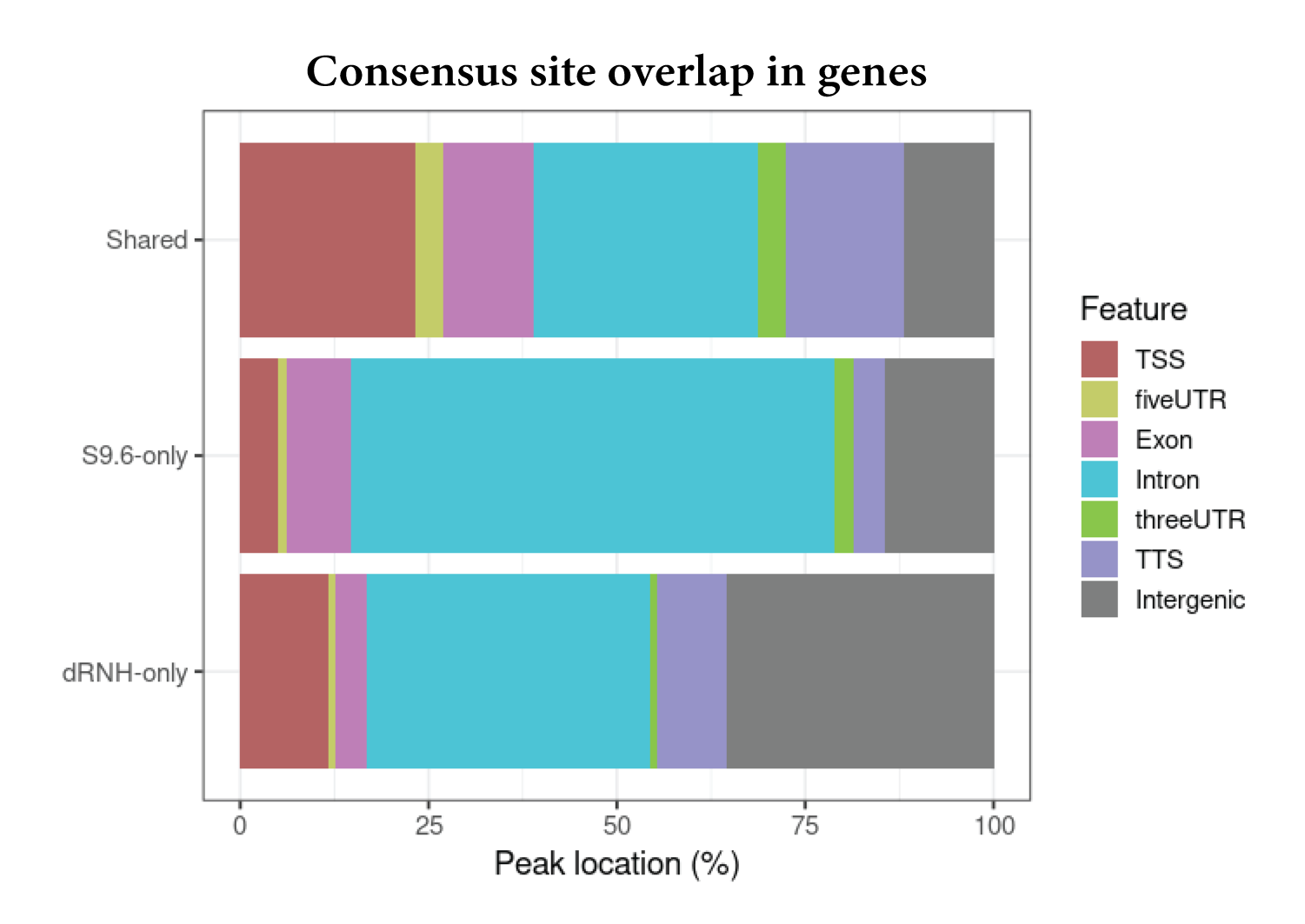

**Figure S5. Annotation plot showing the percent of 500bp summitted consensus sites overlapping with transcriptomic features.** Sites are split by “Shared” (found by S9.6 and dRNH), “dRNH-only”, and “S9.6-only”.

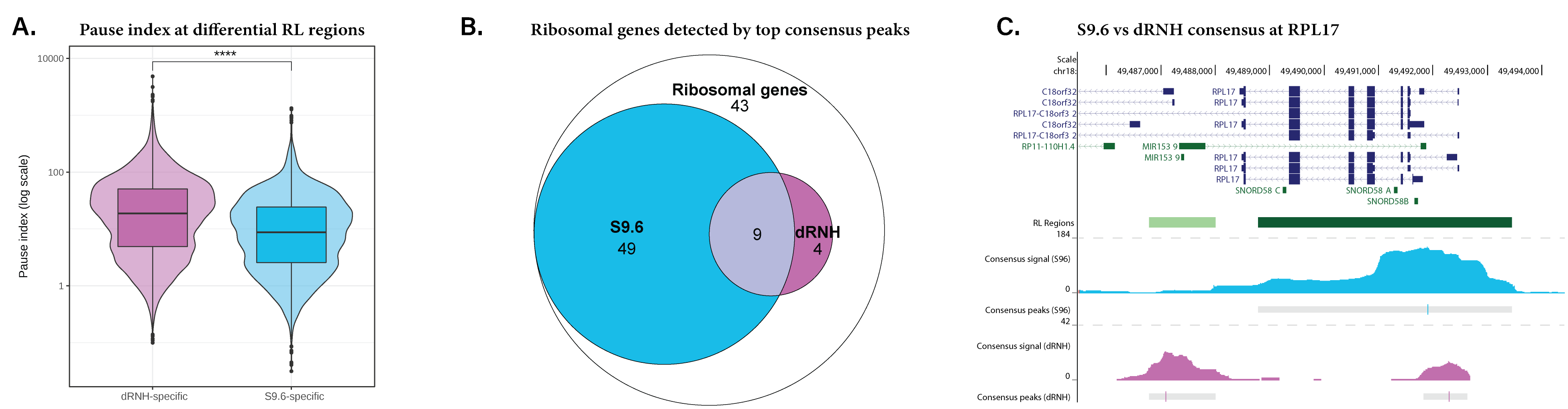

**Figure S6. Extension of Figure 5 results.** (A) A box/violin plot showing the pause index of differential RL regions specific to dRNH and S9.6. Test of means via Wilcoxon rank sum test. (B) An Euler diagram showing the number of genes from the “KEGG Ribosome” pathway overlapping with the top 2000 peaks from S9.6 and dRNH. (C) A UCSC Genome Browser image showing a ribosomal gene (RPL17) which showed increased consensus signal in S9.6 compared to dRNH from the analysis in (A). **** - p < .0001; *** - p < .001; ** - p < .01; * - p < .05; ns – p >= .05.

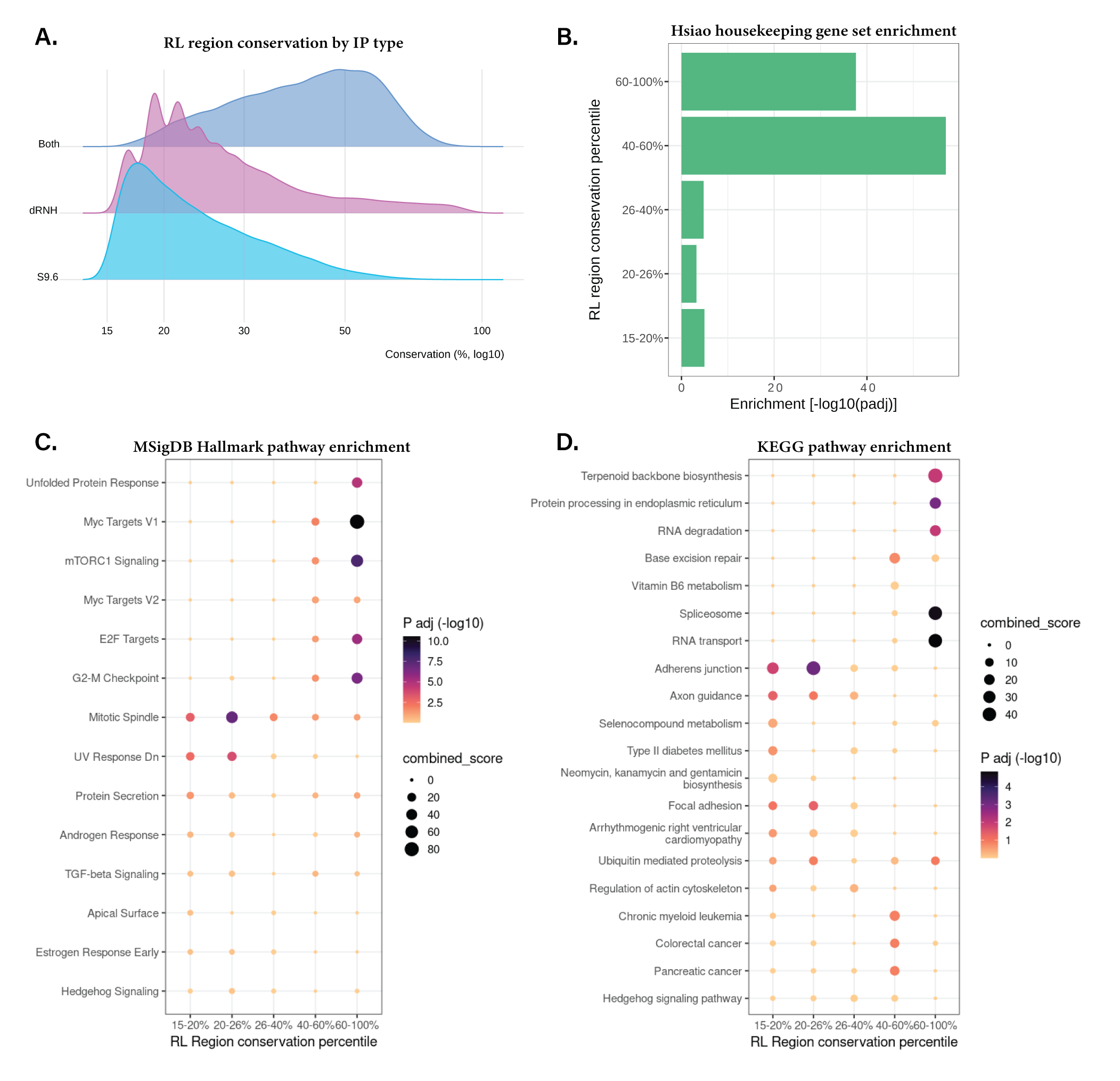

**Figure S7. Extension of Figure 6.** (A) Ridge plot showing the distribution of conservation percentages among the RL regions discovered by dRNH samples, S9.6 samples, or both. (B) Bar plot showing the enrichment of genes overlapping RL regions from each conservation bin within the “HSIAO_HOUSEKEEPING_GENES” gene set (MSigDB C2 collection). P value is from the hypergeometric test with Benjamini Hochberg correction for multiple testing. (C) Pathway enrichment plot showing the significance (via P adjusted value) and effect size (via Combined Score) of MSigDB Hallmark pathway enrichment from the genes overlapping RL regions within each percentile. (D) Same as (C) except with KEGG pathways.
